# Interactions between the myosin Dachs, the adaptor Dlish, and the palmitoyltransferase Approximated mediate Fat-Dachsous signaling

**DOI:** 10.64898/2026.04.14.718308

**Authors:** Xing Wang, Yifei Zhang, Junjie Zhai, Xiaoyi Yang, Seth S. Blair

## Abstract

Bidirectional signaling mediated by heterophilic binding between the giant Drosophila protocadherins Fat and Dachsous (Ds) limits growth through the Hippo pathway and, via patterned binding and cell-by-cell polarization, influences planar cell polarity (PCP) and fate choices along major tissue axes. Prior work showed that the signal transduction initiated by the intracellular domains (ICDs) of Fat and Ds changes the localization and levels of three unusual binding partners: the type XX atypical myosin Dachs, the SH3 domain containing adaptor Dlish, and the DHHC palmitoyltransferase Approximated (App). However, the complex interactions between the three proteins and Fat and Ds are less well understood. Our evidence shows that these proteins play overlapping but distinct roles regulating each other and mediating signaling. Dlish localization and activity requires App binding and palmitoylation. Palmitoylated Dlish helps tether Dachs to the cortex but is also needed to couple Dachs to stabilization by the Ds ICD and destabilization by the Fat ICD. In contrast, Dachs does not localize Dlish but protects it from degradation. Our results also indicate that Fat does not act by changing App levels or localization. We discuss an alternative model based on Dlish’s proposed role as an adaptor for E3 ubiquitin ligases.

## Introduction

Loss of Fat and, to a lesser extent, Ds, causes massive overgrowth of the imaginal disc epithelia, which proliferate during larval stages and during metamorphosis give rise to the appendages and body ectoderm of the adult head and thorax. Loss of Fat or Ds also disrupts the PCP of cell divisions, hairs, and differentiation choices in imaginal and other epithelial tissues, and mispositions tissues such as the crossveins of the wing. Fat and Ds are not only necessary but instructive: in several tissues the strength and polarity of Fat/Ds binding is regulated by a gradient of Ds expression and an opposite gradient of Four-jointed, a Golgi-resident kinase that alters Fat-Ds binding by phosphorylation of their cadherin domains, leading to the cell-by-cell polarization of Fat and Ds to opposite cell faces. Increasing Fat/Ds binding reduces growth, while artificial boundaries and gradients of Fat, Ds or Fj induce overgrowth, alter gene expression, and redirect PCP^1-4^.

Some of these functions are shared by mammalian homologs of Fat and Ds during neurogenesis, proliferation, and different types of PCP^5-9^. Mutations in humans’ Fat homolog FAT4 or the Ds-like protein DCHS1 cause the multisystem defects of Hennekam and Van Maldergem syndromes and have been linked to tumor formation^8,10-12^. Uncovering the mechanisms behind these defects, and how Fat and Ds are used during normal development, is a high priority.

While Fat and Ds can alter cell adhesion and tension^13^, we and others showed that much of Fat and Ds activity is mediated by their intracellular domains (ICDs). As these lack the enzymatic or protein binding domains of typical signal receptors, researchers have used both genetics and binding screens to find components that transduce Fat/Ds signaling. Recent work has focused on the myosin Dachs^14^, the adaptor protein Dlish (also known as Vamana)^15,16^, and the palmitoyltransferase App, a homolog of yeast Erf2 and mammalian ZDHHC8, DCHHC14, and ZDHHC18^17^. These proteins can bind the ICDs of Fat and Ds, and removal of any one of them largely blocks the overgrowth and improves the PCP defects of null *fat* and *ds* mutations.

Previous studies strongly suggest that Fat and Ds ICDs work by changing the levels, localization, and activity of these proteins in the subapical cell cortex, providing a link between Fat-Ds junctions and subapically concentrated components of the Hippo and PCP pathways^2,4^ (Fig. 1). The myosin Dachs induces growth by changing the conformation of, and thereby inhibiting, Warts, the terminal kinase in the Hippo pathway^14,18-20^. Dachs, along with the Ds ICD, also binds and helps concentrate the Spiny-legs isoform of the Fz-Vang PCP pathway protein Prickle^21,22^ and changes membrane tension and the mechanics of tissue polarization^13^. Dlish promotes growth by binding directly to the FERM domain scaffolding protein Expanded (Ex) and reducing its levels, likely by stimulating Ex ubiquitination^23^. Ex normally inhibits growth by increasing Hippo and Warts phosphorylation^24-29^ and by binding and inhibiting Yorkie (Yki), a Yap/Taz family transcriptional co-factor that provides the main growth-inducing output of the Hippo pathway^30-32^.

**Figure 1.**
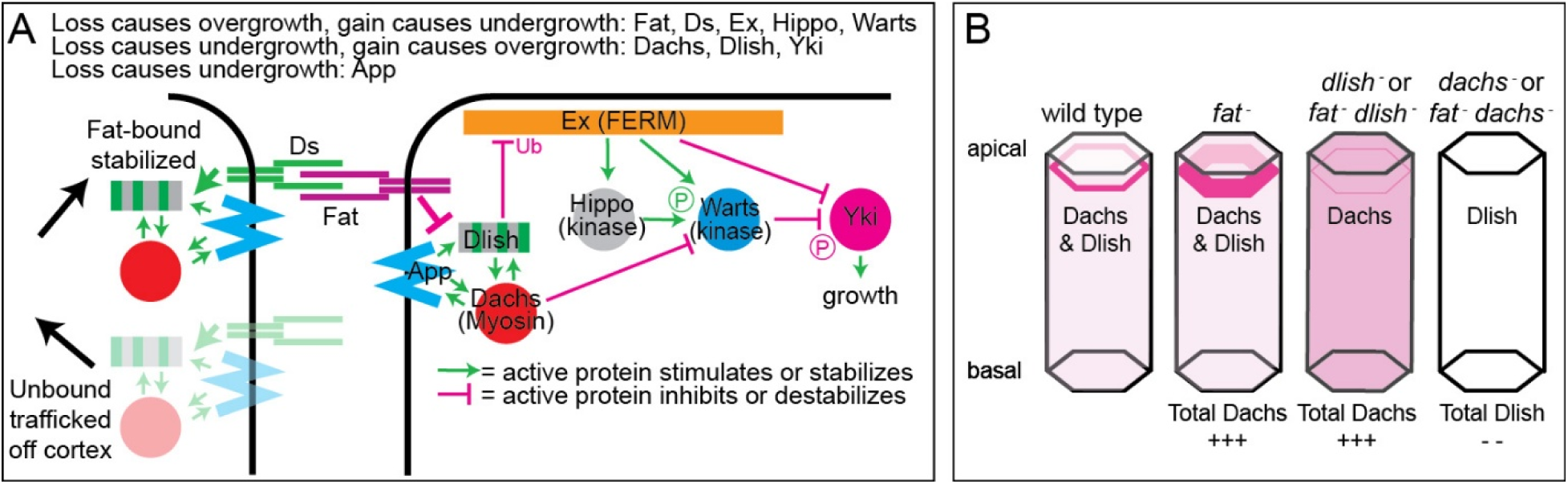
Activity of Dachs, Dlish, and App in the Fat-Ds and Hippo pathways. A. Simplified model of the effects of Fat and Ds on Dachs, Dlish, and App levels and activity, and the effects of Dachs and Dlish on elements of the Hippo pathway. B. Effects of *fat*, *dlish* and *dachs* mutations on the subapical cortical, cytoplasmic, and total levels of Dachs and Dlish proteins in imaginal discs (from refs^14,15,33^ and this study).

The Fat and Ds ICDs have strong effects on the subapical levels of Dachs, Dlish, and App. The Ds ICD may accomplish this by simple binding. However, whether this binding increases or decreases proteins in the subapical cortex may depend on whether cortical Ds is stabilized by forming extracellular junctions with Fat^33^. Fat-bound, junctional Ds would stabilize cortical Dachs, Dlish, and App, but loss of *fat* increases trafficking of Ds and its binding partners away from the cell membrane^33,34^. Blocking this trafficking by the removal of Ds may explain the increased cortical Dachs, Dlish, and App and the greater overgrowth in *fat ds* double mutants compared with *fat* mutants.

In contrast, while Dachs, Dlish, and App can bind to the Fat ICD in vitro, in vivo both Ds-bound and unbound Fat strongly reduce Dachs and Dlish levels at the subapical cell cortex (Fig. 1B), while more weakly reducing App^14-16,33^. The increased cortical Dachs and Dlish is critical for the *fat* mutant overgrowth phenotype. Overgrowth is rescued by loss of Dachs or Dlish, and overexpression of either can mimic *fat* mutant overgrowth^14-16^. Changes in Dlish levels also likely account for Fat’s effects on subapical Ex: Ex directly binds both Dlish^23^ and the Fat ICD^35^, but while subapical Ex is reduced in *fat* mutants^36-39^, the mutant effect is blocked by loss of *dlish*^23^.

How does Fat mediate these effects, how are Dachs and Dlish tethered to the cortex in the absence of Fat and Ds, and what is the role of the palmitoyltransferase App in this process? While App contributes to palmitoylation of the Fat ICD, this does not explain App’s activity in null *fat* mutants^17,40^. Importantly, Dlish and Dachs are dependent on each other (Fig. 1B) and on App for their concentration in the subapical cortex, even in the absence of *fat*; conversely, increases in Dachs increase cortical Dlish and vice-versa^15,16^. The second of Dlish’s three SH3 domains binds directly to an SH3 binding motif in the C-terminus of Dachs, Dlish can also bind App, and our in vitro evidence suggested that App palmitoylates Dlish. We therefore hypothesized that 1) App-palmitoylated Dlish helped tether the Dachs-Dlish complex to the subapical plasma membrane, and 2) inhibition of Dlish palmitoylation by the Fat ICD provided a mechanism by which Fat regulates the cortical levels of the complex^15^.

However, others have questioned the palmitoylation data and thus the putative role for both Dlish and App^16,41^, and previous studies left largely untested the instructive or permissive roles for App and other complex members.

Here, we provide strong evidence that direct interactions between Dlish and Dachs are necessary for their function and localization in vivo. We further show that Dlish is palmitoylated, that Dlish lipidation is needed to properly concentrate Dachs in the cell cortex, and that the normal localization of Dlish and Dachs requires binding between Dlish and App. Our evidence also argues that Dlish not only helps localize Dachs but also couples Dachs to Fat or a mediator of Fat activity. However, we also show that App levels are not limiting, suggesting that Fat-regulated changes in App levels or availability are unlikely to mediate the *fat* mutant phenotype.

Intriguingly, we also provide new evidence that the system controls not simply the localization of protein partners but total protein levels in the cell. Previous work showed that loss of *fat* or *dlish* increased the total levels of Dachs per cell, in *fat* mutants chiefly in the cortex, in *dlish* mutants chiefly in the cytoplasm^15,42^ (Fig. 1B). Here we show that the major effect of losing *dachs* is not to untether Dlish from subapical cortex but rather to reduce Dlish levels throughout the cell. Given previous evidence that Dlish couples Ex to an E3 ubiquitin ligase^23^, we discuss models that incorporate these findings.

## Results

### Normal Dachs localization and activity requires direct binding to Dlish

The simplest working hypothesis is that palmitoylated Dlish acts by localizing Dachs, recruiting Dachs to cortical membranes in the subapical region of imaginal disc epithelial cells. Once at the subapical cortex Dachs inhibits the Hippo pathway component Warts, while Dlish reduces the levels of Ex.

Purified Dachs and Dlish can directly bind each other in vitro and that binding requires Dlish’s second SH3 domain and a specific SH3 binding motif in the Dachs C-terminus^15,16^. The evidence to date suggests that this direct binding mediates Dlish’s effects on Dachs: in vivo, a Dlish lacking the second SH3 domain (DlishΔSH3-2) does not rescue the Dachs localization defects caused by RNAi knockdown of *dlish^16^*.

As a further test, we examined the in vivo activity and subcellular localization of a Dachs construct lacking its C-terminal SH3-binding motif (DachsΔSH3bd). In wing discs, overexpression of wild type Dachs in the posterior of wing discs using *hh-gal4* caused overgrowth and increased expression of the Yki target *ban3-GFP* (Fig. 2B-C’). In contrast, expression of DachsΔSH3bd in the posterior of the wing caused no obvious overgrowth and only a very weak effect on *ban3-GFP* (Fig. 2E-F’). As expected, this correlated with a difference in Dachs localization: overexpressed Dachs preferentially concentrated in the subapical cell cortex (Fig. 2C”), while DachsΔSH3bd localized more diffusely throughout the cell (Fig. 2F”).

**Figure 2.**
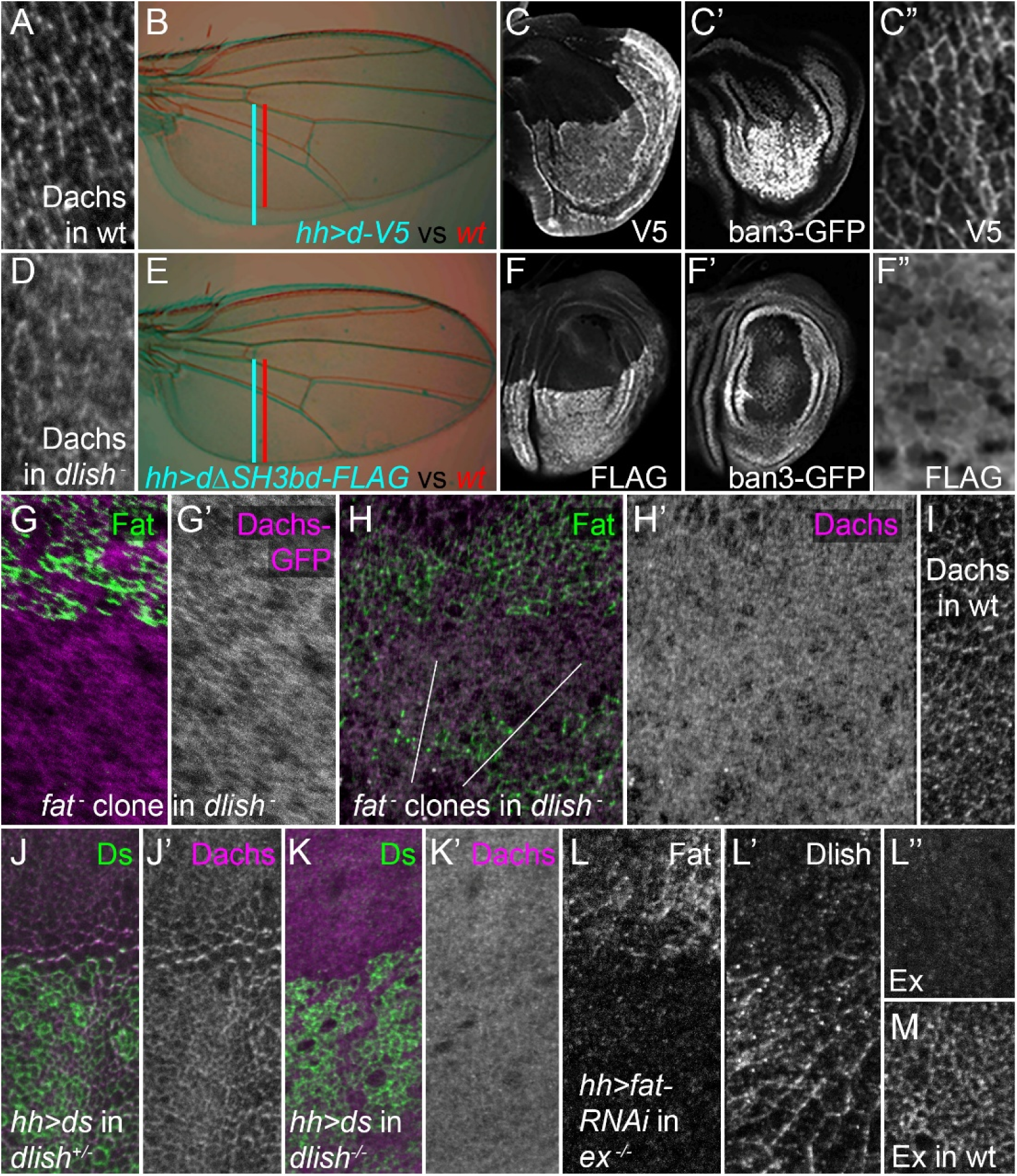
Effects of *dlish*, *fat*, *ds* on Dachs and *ex* on Dlish. A,D. Anti-Dachs staining in wild type (wt, A) and *dlish^B1601^* homozygous mutant (D) wing discs. B-C’, E-F’. Different effects of posterior, *hh-gal4*-driven expression of *UAS-d-V5* (B-C”) and *UAS-dΔSH3bd-FLAG* (E-F”). B,E. Adult wings and posterior growth in experimental (blue) and wt (red); blue and red bars show posterior growth. C-C’, F-F”. Wing discs showing posterior growth and tagged Dachs distribution by staining with anti-V5 (C,C”) or anti-FLAG (F,F”), and their effects on the Yki target *ban3-GFP* (C’,F’). G-H’. Dachs-GFP (magenta, grey G,G’) and anti-Dachs staining (magenta, grey, H-H’) in *dlish^B1601^*/*dlish^Y003^* discs containing *fat^fd^* homozygous clones marked by the absence of GFP (green). I. Anti-Dachs staining in *fat^fd^ dlish^Y003^/+* disc. J-K’. Anti-Dachs staining (magenta, grey) after posterior, *hh-gal4*-driven expression of UAS-Ds (J,K, anti-Ds in green) in *dlish^B1601^* heterozygote (J,J’) and *dlish^B1601^* homozygous (K,K’) wing discs. L-L’’. Anti-Fat (L) and anti-Dlish staining (L’) after posterior, *hh-gal4-*driven expression of *UAS-fat-RNAi* (BSC 29566) in *ex^e1^* homozygous wing disc, and anti-Ex staining (L”) compared with wt (M). Compared disc pairs (A vs D, H’ vs I, L” vs M) were fixed and stained in the same well, mounted on the same slide, and images captured and adjusted with identical settings.

### Dlish is required for the effects of Fat on cortical Dachs

Interestingly, despite the more diffuse distribution of DachsΔSH3bd compared with wild type Dachs, there is still residual concentration of DachsΔSH3bd at the subapical cell cortex (Fig. 2F”). Similarly, we often detected residual subapical cortical Dachs in homozygous *dlish* null wing discs (Fig. 2A,D). If the sole role of Dlish was to increase Dachs at subapical membranes, this simple localization model would predict that the residual cortical Dachs present in *dlish* null mutant discs would still be capable of inhibiting Warts and Ex, thereby increasing tissue growth. This prediction is supported, as *dlish* null mutations have weaker undergrowth than that caused by loss of *dachs*, and *dachs* overexpression can weakly increase Yki activity in the absence of *dlish*^15^.

However, the simple localization model would also predict that the residual cortical Dachs present in *dlish* mutants would still be sensitive to loss of Fat. Instead, we found that *fat* null clones did not detectably increase cortical Dachs in a *dlish* null mutant background, as assessed using either a Dachs-GFP transgene (Fig. 2G,G’) or anti-Dachs staining (Fig. 2H,H’). Moreover, the higher level of diffuse cytoplasmic Dachs found in *dlish* mutants was also unaffected by *fat* mutant clones; compare Fig. 2H’ to wild type (wt) in Fig. 2I. This agrees with independent findings of Matakatsu and Fehon^33^, and indicates that Dlish not only helps concentrate Dachs in the subapical cortex but also links subapical Dachs to Fat function.

### Dlish is required for the effects of Ds on cortical Dachs

Subapical Dachs is also sensitive to the ICD of Ds in vivo, and our results indicate that these effects also depend on Dlish. When cortical Ds is stabilized, for instance by binding to the Fat ECD, the Ds ICD increases cortical Dachs^13,33^. Overexpressed Ds can increase cortical Dachs, especially at boundaries between normal and Ds-overexpressing cells^43-46^. Within the region of Ds overexpression competition for binding to endogenous Fat can limit the levels of junctional Ds, but at boundaries the overexpressed Ds attracts and caps the Fat on the adjacent normal cells, increasing the stability of both Ds and Fat at boundary cell faces. The capping also depletes Fat on the non-boundary cell faces of normal cells (Fig. S1C), which in turn encourages endogenous Ds in those cells to bind to the Fat in adjacent non-boundary cells, polarizing Ds-bound Dachs; this effect can propagate over several cells^45,46^ (Fig. 2J,J’). Removal of Dlish blocked the accumulation and polarization of Dachs at boundary and adjacent cell faces (Fig. 2K,K’). This strongly suggests that the stabilization of Dachs by the Ds ICD depends on the formation of a Dachs/Dlish complex.

The increased Dachs at Ds overexpression boundaries is also thought to account for the boundary-specific expression of Yki targets: the boundary-specific Yki activity is blocked by loss of Dachs^43,44^. As expected, the activation of the Yki target *ban3-GFP* at boundaries of *hh-gal4*-driven *UAS-ds* overexpression was also blocked by loss of Dlish (Fig. S1A,B).

We note that reliance of Ds-Dachs interactions on Dlish is also consistent with a complementary result of Matakatsu and Fehon^33^. When the turnover of cortical Ds is increased by loss of Fat binding this destabilizes cortical Dachs and Dlish, likely via trafficking of a Ds/Dlish/Dachs complex away from the membrane; the destabilization of cortical Dachs depends on Dlish.

### Fat’s effects on Dlish are not bridged by Expanded

If Dlish activity extends beyond simple cortical localization of Dachs, one possible route is through interactions with Fat: Dlish binds Dachs-regulating portions of the Fat ICD in vitro^15,16,33^ and this binding may mediate Fat’s effects on Dachs. However, recent studies suggest that the binding between the Fat ICD and Dlish might not be direct, and instead might be bridged by Ex. We previously showed that Dlish can directly bind Ex^23^, and others found that bacterially produced Ex, but not the equivalent Dlish, can directly bind the Fat ICD^35^. We therefore tested whether Ex acts as a functional bridge between Fat and Dlish that mediates Fat’s effects on Dlish localization and levels. However, we found that *fat* knockdown still increased cortical Dlish in an *ex* null background (Fig. 2J,K). We also note that *ex* mutant overgrowth, unlike *fat* mutant overgrowth, does not act by changing the levels of subapical Dlish or Dachs^23^.

An Ex-independent effect of Fat on Dlish and Dachs may explain why *fat* loss further increases overgrowth of *ex* null mutant discs^39^. Nonetheless, the Ex and Fat/Dlish pathways are not completely independent, as loss of *fat* reduces and loss of Dlish increases Ex levels in vivo^23,36-39^. In vitro Dlish can also immmuno-precipitate (IP) the Slimb ubiquitin ligase, which is known to target Ex, and Dlish increases Ex ubiquitination; we proposed that Fat binds and sequesters or inhibits Dlish and protects Ex from Dlish-aided ubiquitination^15,23^.

### Dachs stabilizes but does not localize Dlish

The results above suggest that Fat destabilizes subapical Dachs by binding to and regulating Dlish. However, this is an oversimplification as the interactions between Dlish and Dachs are in some sense reciprocal: Dachs loss strongly reduces cortical Dlish in both wild type, *fat* mutant^15,16^ and *fat ds* double mutant backgrounds^33^. In fact, a strong RNAi knockdown of *fat* that increased cortical Dlish levels in wild type discs did not detectably increase cortical anti-Dlish staining in a *dachs* null background (Fig. 3D-E’).

**Figure 3.**
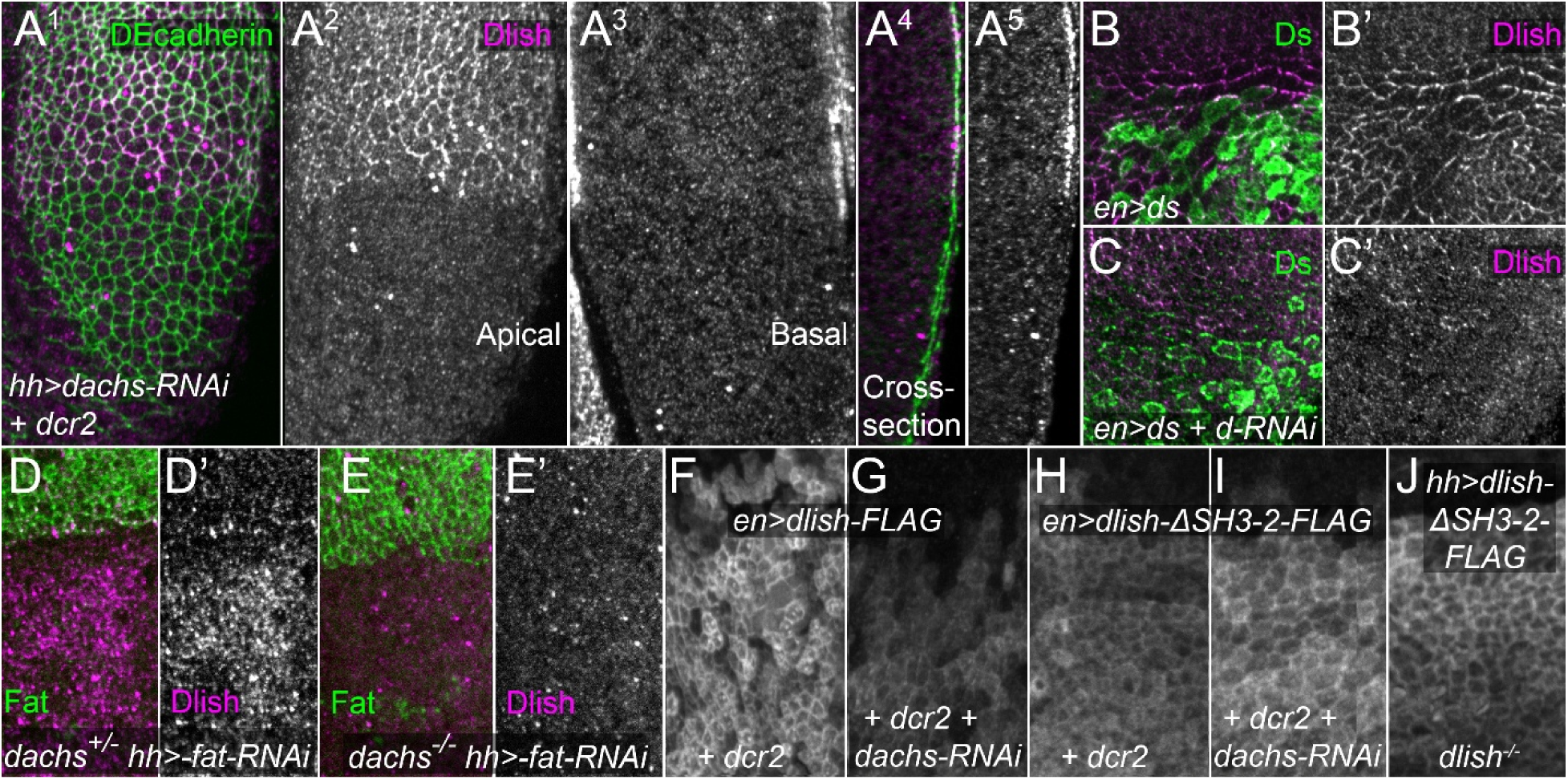
Effects of *dachs* knockdown on Dlish. A^1^-A^5^. Effects of posterior, *hh-gal4*-driven *UAS-dcr2* and *UAS-dachs-RNAi* on subapical cortical (A^1^,A^2^) and basolateral cytoplasmic (A^3^) anti-Dlish staining (magenta, grey); anti-DEcadherin counterstain in green. Cross-sections shown in A^4^,A^5^, with apical to the right. B-C’. Anti-Dlish staining (magenta, grey) after posterior, *en-gal4*-driven expression of *UAS-ds* (anti-Ds in green) without (B,B’) or with (C,C’) *UAS-dachs-RNAi*. D-E’. Anti-Dlish staining (magenta, grey) after posterior, *hh-gal4*-driven expression of *fat-RNAi* in *dachs^210^*/+ (D,D’) versus *dachs^210^*/*dachs^CG13^* (E,E’). F-J. Anti-FLAG staining after posterior, *en-gal4*-driven expression of *UAS-dcr2* and either *UAS-dlish-FLAG* (F,G) or *UAS-dlishΔSH3-2-FLAG* (H,I) without (F,H) or with (G,I) expression of *UAS-dachs-RNAi*, or (J) after posterior, *hh-gal4*-driven expression of *UAS-dlishΔSH3-2-FLAG* in a *dlish^B1601^* homozygote.

We originally hypothesized that, as with Dlish’s effect on Dachs, Dachs’ effect on Dlish was in part through recruitment to the subapical cortex. While the myosin motor properties of Dachs are in question, Dachs can bind actin^47^, and thus we proposed that Dachs tethered Dlish to the cortical cytoskeleton^15^.

However, our new results suggest that Dachs regulates Dlish protein levels but not its localization. Notably, the effect of Dachs on Dlish differs from the effect of Dlish on Dachs. After loss or knockdown of Dlish, cortical Dachs is reduced but cytoplasmic levels increase, consistent with an effect on Dachs localization^15,16^ (Fig. 2A,D). In contrast, *dachs* knockdown decreased the levels of both subapical and cytoplasmic Dlish (Fig. 3A^1^-A^5^); the effect in *dachs* null mutant wing discs is extremely strong (Fig. S2A^3^,B^3^,C^3^).

These findings suggest that Dachs directly or indirectly protects Dlish from degradation. This hypothesis is supported by two additional results. First, overexpressed Dlish-FLAG still preferentially concentrates in the subapical cortex even after strong RNAi knockdown of *dachs* (Fig. 3F,G). Second, overexpressed Dlish-ΔSH3-2-FLAG, which lacks the second SH3 domain required for binding Dachs in vitro^15,16^, is also subapically concentrated^16^, even in wing discs lacking endogenous Dlish (Fig. 3J), and is not detectably reduced by strong *dachs* knockdown (Fig. 3H,I).

The mechanism by which Dachs stabilizes Dlish is as yet uncertain. Reducing the activity of two ubiquitin ligases known to reduce subapical Dlish or Dachs, Fbxl7 and Early girl (Elgi)^15,41,48,49^ did not detectably increase the levels of endogenous Dlish after *dachs* loss or knockdown (Fig. S2).

### App levels are not limiting

Previous studies indicated that the activity of the DHHC palmitoyltransferase App helps recruit or stabilize the Dlish-Dachs complex to the subapical cortex. Loss of *app* reduces subapical and increases cytoplasmic Dachs in both wild type and *fat* mutant cells^15,17,40^ (Fig. 4A-C”). Similarly, a strong *UAS-app-RNAi* from the VDRC RNAi project^50^ reduced cortical and increased cytoplasmic Dlish (Fig. 4D,D’) and gave the crossvein spacing defect typical of *app*, *dlish*, and *dachs* mutants (Fig. 4H). Alternative splicing of the *app* transcript is predicted to produce four isoforms of App that have the DHHC domain required for palmitoyltransferase activity but differing in their C-terminal tails (Fig. S3). The VDRC *app-RNAi* targets only the *app-RJ/RK* mRNAs, indicating that the App-PJ/PK protein is a critical isoform in the wing.

**Figure 4.**
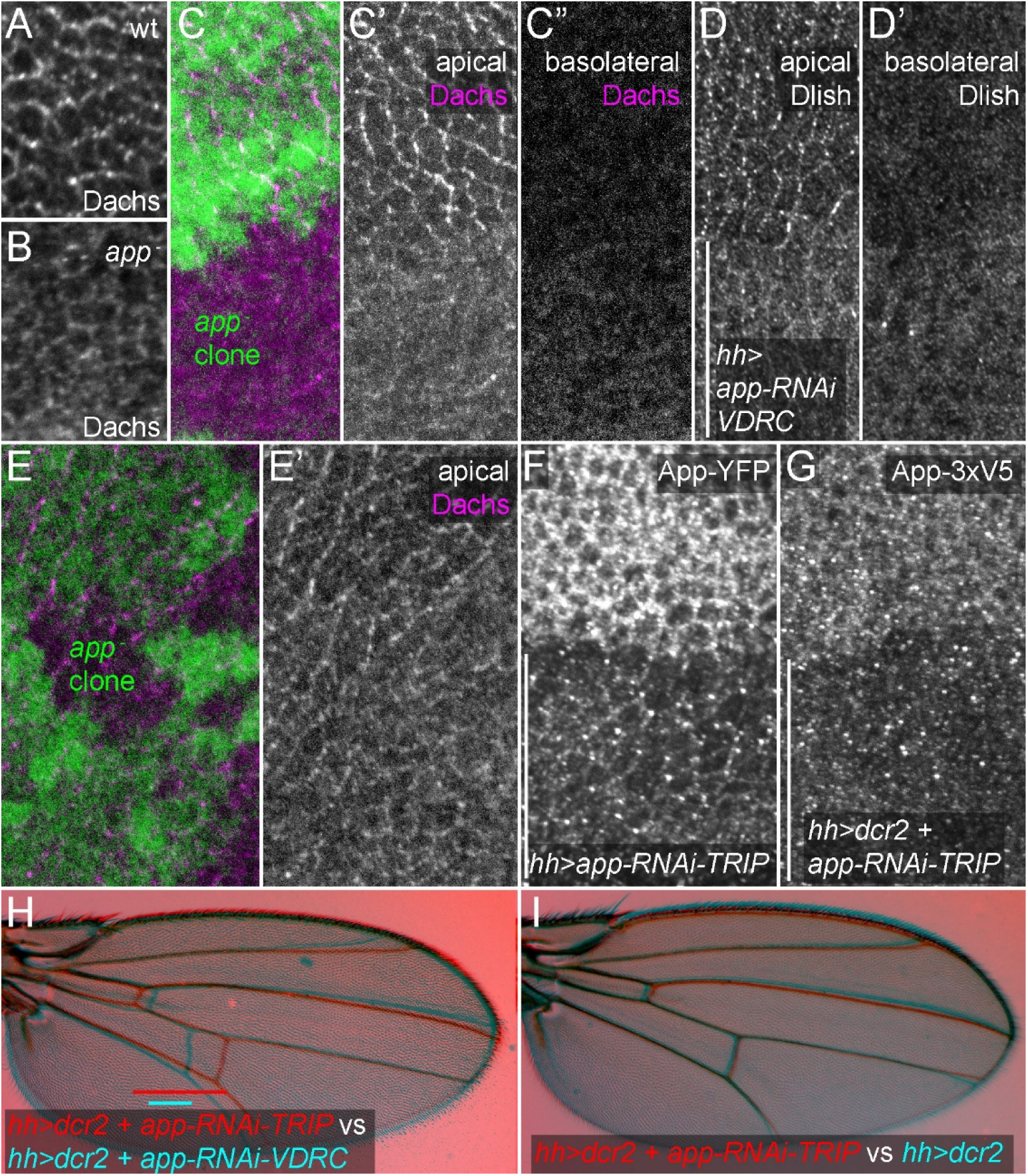
Effects of *app* loss or knockdown. A.B. Anti-Dachs staining in wt (A) and *app^12-3^*/*Df(3L)BSC730* (B); wings discs fixed and stained in the same well, mounted on the same slide, and images captured and adjusted with identical settings. C-C” Effect of large *app^12-3^* homozygous mutant clone, identified by absence of GFP (green), on anti-Dachs staining (magenta, grey) at apical (C,C’) and basolateral (C”) focal planes. D-D’. Effect of posterior (identified with bar), *hh-gal4*-driven expression of the strong VDRC *UAS-app-RNAi*, on anti-Dlish staining at apical (D) and basolateral (D’) focal planes. E,E’. Effect of small *app^12-3^* homozygous mutant clone, identified by absence of GFP (green), on apical anti-Dachs staining (magenta, grey). F,G. Effect of posterior, *hh-gal4*-driven expression of weak *TRIP UAS-app-RNAi* on endogenous tagged *app^YFP^* (F) or on *app^3xV5^* with *UAS-dcr2* (G). H,I. Comparison of adult wing crossvein spacing (bars) after posterior *hh-gal4*-driven expression of *UAS-dcr2* and the weak TRIP *UAS-app-RNAi* (red) to either the strong VDRC *UAS app-RNAi* (H, cyan) or the UAS-drc2 control (I, cyan).

As noted above we originally proposed that App provides a route through which the Fat and Ds ICDs might regulate the Dlish/Dachs complex, as in vitro App can bind the Fat and Ds ICDs as well as Dachs and Dlish^15,17,33,40^. In vivo much of the App concentrates in the subapical cortex^17,33^ (Fig. 4F,G) and it is possible that changes in subapical levels controls Dlish palmitoylation and thus Dlish and Dachs activity. Loss of *fat* and *ds* can increase App in the subapical cortex, although the increase is weaker than the increase of Dachs and Dlish and may depend on the increase in Dachs; conversely, App can by increased by increasing junctional Ds or Dachs and is at lower levels on the Fat side of Fat-Ds junctions^33^.

However, even very low levels of App have sufficient enzymatic activity to control Dachs and Dlish. The amount of App that perdures in small *app* null clones is able to normally localize Dachs (Fig. 4E,E’). A weaker *UAS-app-RNAi* line from the TRiP RNAi project^51^ that targets all of the DHHC-containing splice isoforms of App (Fig. S3) greatly reduces the levels of endogenous, epitope-tagged App (Fig. 4F,G) but has no detectable effect on crossvein spacing, the most sensitive marker of App activity (Fig. 4H,I). Thus, App levels are not limiting. Either App-mediated palmitoylation is largely permissive, not instructive, or any regulation of App activity by Fat or Ds involves unknown, limiting co-factors.

### Dlish can be palmitoylated at multiple cysteines

The changes in Dlish localization after App knockdown do not resemble the effect of Dachs knockdown, as the former increases and the latter decreases cytoplasmic Dlish (compare Fig. 3A^3^ to Fig. 4C”,D’). This suggests that App is not directly regulating Dachs activity, consistent with the lack of detectable Dachs palmitoylation in S2 cells^15,40^. Instead, in vitro a fraction of Dlish appears to be palmitoylated as assessed using the Acyl-Biotin exchange (ABE) assay^15^. However, others have questioned whether the biotinylated band shown in our ABE assay is specific for Dlish; moreover, loss of C5 from Dlish, which the CSS-Palm predictor^52^ suggests is the cysteine most likely to be palmitoylated, does not disrupt Dlish function^16,41^.

We therefore repeated the ABE assay on protein isolated from S2 cells with anti-FLAG IP and found that the biotinylated band in the assay is only present in cells expressing Dlish-FLAG (Fig. 5A) and was eliminated by mutating all of Dlish’s potentially palmitoylated cysteines to serine (Dlish-all C to S-FLAG) (Fig. 5B). Next, as the cysteine acylation detected by the ABE assay is not strictly specific for palmitoylation, we confirmed Dlish palmitoylation in S2 cells with a click chemistry assay that uses an azide-linked biotin probe to test whether an alkyne-modified palmitate analog is incorporated into Dlish-FLAG. The probe bound much more strongly to IP’d Dlish-FLAG than to our controls, a Dlish-FLAG treated with HAM to cleave off alkyne-palmitate, or equal amounts of Dlish-all C to S-FLAG (Fig. 5C).

**Figure 5.**
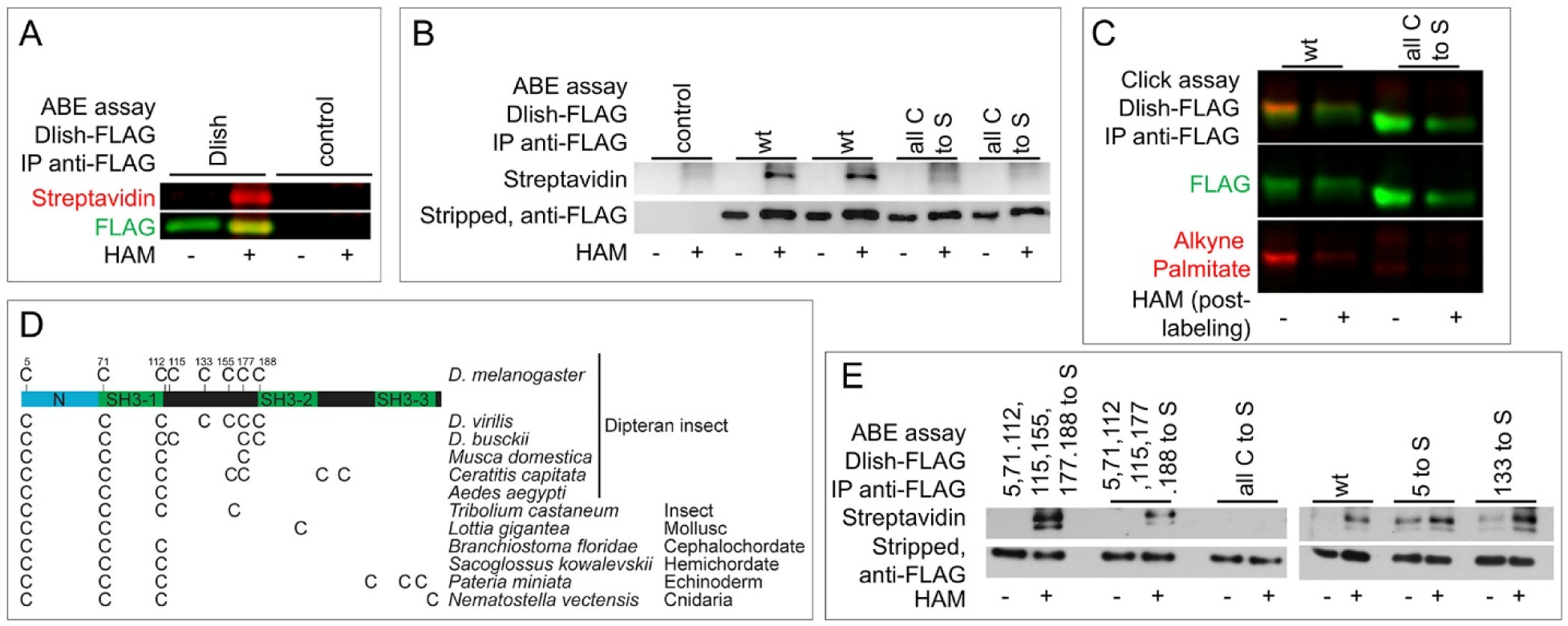
Palmitoylation of Dlish. A. In vitro ABE assay showing that HAM-treating anti-FLAG IP extract labels a band of the correct MW only if cells express Dlish-FLAG. B. In vitro ABE assay comparing labeling of Dlish-FLAG and Dlish-all C to S-FLAG. C. Click it assay comparing incorporation of alkyne-labeled palmitate into Dlish-FLAG and Dlish-all C to S-FLAG in S2 cells, and removal of palmitate by HAM treatment. D. Differences in the numbers and positions of cysteines in Dlish homologs from different invertebrate species. E. In vitro ABE assay assessing palmitoylation of Dlish-FLAG mutants in which selected cysteines were converted to serines.

In order to test the importance of Dlish palmitoylation in vivo we attempted to identify which of its eight cysteines are S-palmitoylated. Predicted Dlish homologs are found in taxa from Cnidarians to Cephalochordates (but apparently not vertebrates) and most of these preserve cysteines homologous to C5, C71 and C112 (Fig. 5D), suggesting they are critical for function. Nonetheless, in contrast to Dlish-all C to S-FLAG (Fig. 5B), we obtained positive ABE assays after converting all except C133 to S, all except 133 and 155 to S, and after mutating either just C133 or C5 to S (Fig. 5E). This suggests that in S2 cells many, and perhaps all, of the cysteines can be palmitoylated, and so we limited our in vivo tests to Dlish-all C to S.

### Dlish lipidation is necessary for function in vivo

Overexpression of wild type Dlish-FLAG in the posterior of the wing with *hh-gal4* causes substantial overgrowth (Fig. 6A,B); anti-Dlish staining is more highly concentrated at the subapical cell cortex (Fig. 6E) and accompanied by increased subapical Dachs^15,16^ (Fig. 6E’,E^x^). In contrast, Dlish-all C to S-FLAG did not cause overgrowth (Fig. 6C), concentrated in the cytoplasm (Fig. 6F), and increased cytoplasmic Dachs (Fig. 6F’,F^x^). This suggests that Dlish-all C to S-FLAG still bound Dachs but no longer tethered Dachs to the subapical cell membrane.

**Figure 6.**
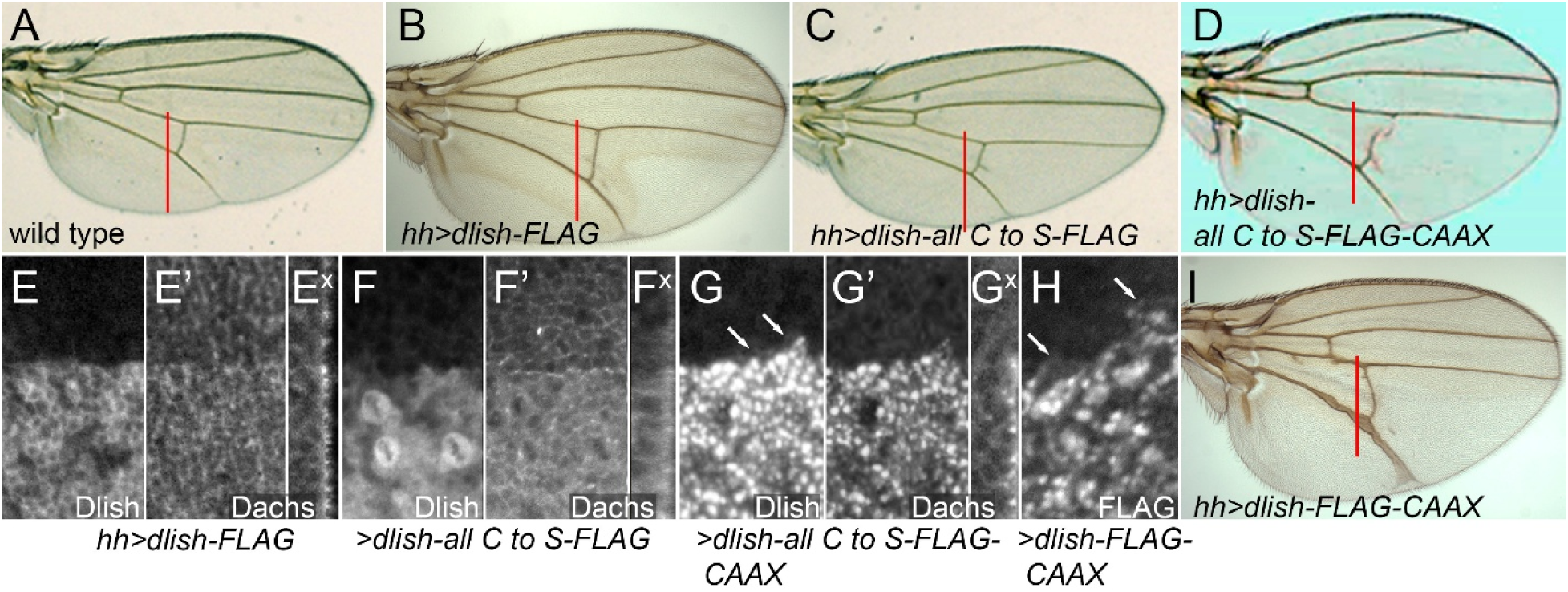
In vivo effects of removing and adding lipidation targets in Dlish. A-E,I. Comparison of posterior wing growth in wt (A, posterior growth shown by red bar) and wings with posterior, *hh-gal4* driven expression of *UAS-dlish-FLAG* (B), *UAS-dlish-all C to S-FLAG* (C), *UAS-dlish-all C to S-FLAG-CAAX* (D), and *UAS-dlish-FLAG-CAAX* (I). E-H. Third instar wing discs with posterior expression of *UAS-dlish-FLAG* (E-E^x^), *UAS-dlish-all C to S-FLAG* (F-F^x^), and *UAS-dlish-all C to S-FLAG-CAAX* (G-G^x^). E,F, and G show anti-Dlish; E’, F’, and G’ show apical anti-Dachs; E^x^,F^x^, and G^x^ show cross-sections of anti-Dachs with apical to the right. H. Anti-FLAG staining after *hh-gal4*-driven expression of *UAS-dlish-CAAX*. Arrows in G and H point to staining in apical cell membranes.

It is possible that removal of cysteines from Dlish affects Dlish structure and binding independent of its palmitoylation. However, much of Dlish function and binding is thought to be mediated by its SH3 domains, and cysteines are not essential for SH3 structure. Nor does the Alphafold Protein Structure Database predict any disulfide bonds in Dlish^53-55^. To rule out additional roles for the Dlish cysteines we first tested whether their loss affected in vitro binding between Dlish and previously published binding partners^15,23^. HA-tagged Dlish-all C to S still co-IP’d with FLAG-tagged Dachs, the Fat and Ds ICDs (FatΔECD and DsICD), the casein kinase Dco, and the E3 ubiquitin ligase Fbxl7; only the binding to App appeared reduced (Fig. 7A,B).

**Figure 7.**
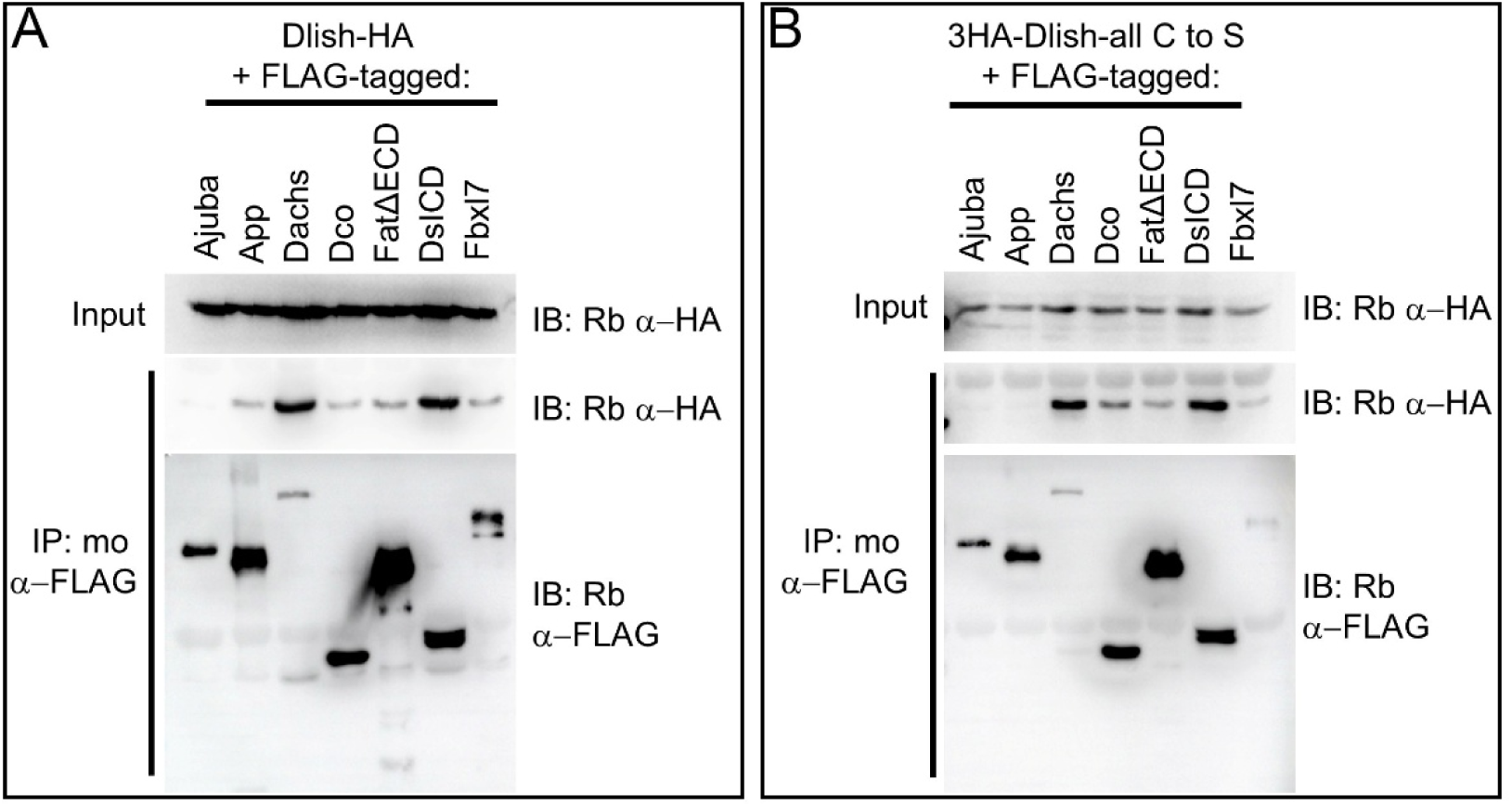
In vitro co-IP assay of binding in S2R+ cells between Dish-HA (A) or 3HA-Dlish-all C to S (B) and FLAG tagged Ajuba, App, Dachs, Dco, FatΔECD, DsICD, and Fbxl7.

To further test whether the altered in vivo activity of Dlish-all C to S-FLAG is due to its lost lipidation we added to it a C-terminal CAAX prenylation motif. The CAAX site is distant from the endogenous Dlish cysteines and is thus unlikely to reproduce any effect endogenous palmitoylation might have on Dlish conformation or binding to specific protein targets. Nonetheless, expressing Dlish all C to S-FLAG-CAAX in the posterior of the wing with *hh-gal4* caused overgrowth that was as strong as that caused by wild type Dlish-FLAG or Dlish- FLAG-CAAX (Fig. 6B,D,I). The similar activities of Dlish, Dlish-CAAX, and Dlish-all C to S-CAAX strongly suggest that Dlish’s cysteines largely act to confer Dlish lipidation and, thereby, membrane localization.

### Dlish-CAAX complexes induce autolysosome-like structures

After *hh-gal4*-driven expression, CAAX-tethered versions of Dlish and Dlish-all-C-to-S were found in the subapical cell cortex (arrows, Figs 6G,H and S4A,A’). However, Dlish-CAAX also concentrated at high levels in very large apical and apico-lateral structures that could occupy as much as half the diameter of the cell. Structures like these have not, to our knowledge, been previously described in imaginal disc cells; they did not-colocalize with the Golgi and were adjacent to but apparently outside of the ER lumen (Fig. S4B,C). Instead, they overlapped with accumulation of a GFP-tagged version of the autolysosomal marker ATG8 (Fig. S4D,D’) and with Lysotracker staining (Fig. S4E,F), which marks acidic cell compartments. Small Lysotracker-positive puncta are found throughout imaginal discs but increased greatly in size in regions expressing Dlish-CAAX (Fig. S4F””). This suggests that aggregated or misfolded Dlish-FLAG-CAAX accumulates in, and increases the size of, autolysosome-like compartments.

Interestingly, these structures also wholly or partially co-localized with atypical concentrations of some of the proteins known to act with Dlish or the Hippo pathway, including Dachs (anti-Dachs in Figs 5G’,G^x^ and S4A”^a,b^; Dachs-GFP in Fig. S4F^3^) and Warts:V5 (Fig. S4G); Ex was weakly affected, although this required the stronger overexpression of Dlish-CAAX driven by *ap-gal4* (Fig. S4H,H’,J,J’). While it is tempting to ascribe these concentrations to the formation of specific complexes with Dlish-CAAX, we favor a more general change in trafficking or autolysosomal activity induced by Dlish-FLAG-CAAX, especially as Dlish-FLAG-CAAX also induced abnormal accumulations of Cad99C (Fig. S4I,I’), an apically concentrated protocadherin not functionally linked to the Fat/Ds or Hippo pathways in the wing^56^.

### Disrupting Dlish-App binding alters Dlish localization and function in vivo

We previously showed that semi-purified App can bind to in vitro translated Dlish^15^. We therefore examined this interaction in more detail to test whether it is important for the cortical localization and activity of Dlish in vivo.

We first investigated the App domains necessary for binding Dlish. Our RNAi evidence above suggests that the alternative splice isoforms producing the App-PJ/K proteins are critical for activity in the wing disc. We found binding between purified, bacterially-produced GST-Dlish-and bacterially-produced MBP-App(1-349) that has the App regions common to all the DHHC-containing App isoforms; however, we also found binding to an MBP-App(350-694) made largely of the C-terminal tail specific to App-PJ/K (Fig. S5). As at least two App domains are sufficient for Dlish binding we did not use this data for in vivo tests.

We then focused on the App-binding domains in Dlish. As summarized in Fig. 8C, we previously showed that the Dlish region spanning the evolutionarily conserved N terminus (NT) and the first SH3 domain was sufficient and necessary for App binding^15^. However, Misra and Irvine found that deleting any single SH3 domain, including SH3-1, did not block subapical concentration of Dlish in vivo^16^, which would be surprising if SH3-1 was necessary for functional interactions between Dlish and App. Our new results suggest that it is the NT, rather than the adjacent SH3-1, that plays the stronger role in App binding: App co-IPs equally well to Dlish-NT+SH3-1, Dlish-NT+SH3-2, and Dlish-NT+SH3-3 (Fig. 8A), and can also co-IP Dlish-NT alone (Fig. 8B). However, while App does not bind to a Dlish construct that lacks NT+SH3-1^15^, we have detected some App binding to Dlish lacking only the NT (Fig. 8B), suggesting either some involvement of the SH3-1 domain or a cooperative effect from all three SH3 domains.

**Figure 8.**
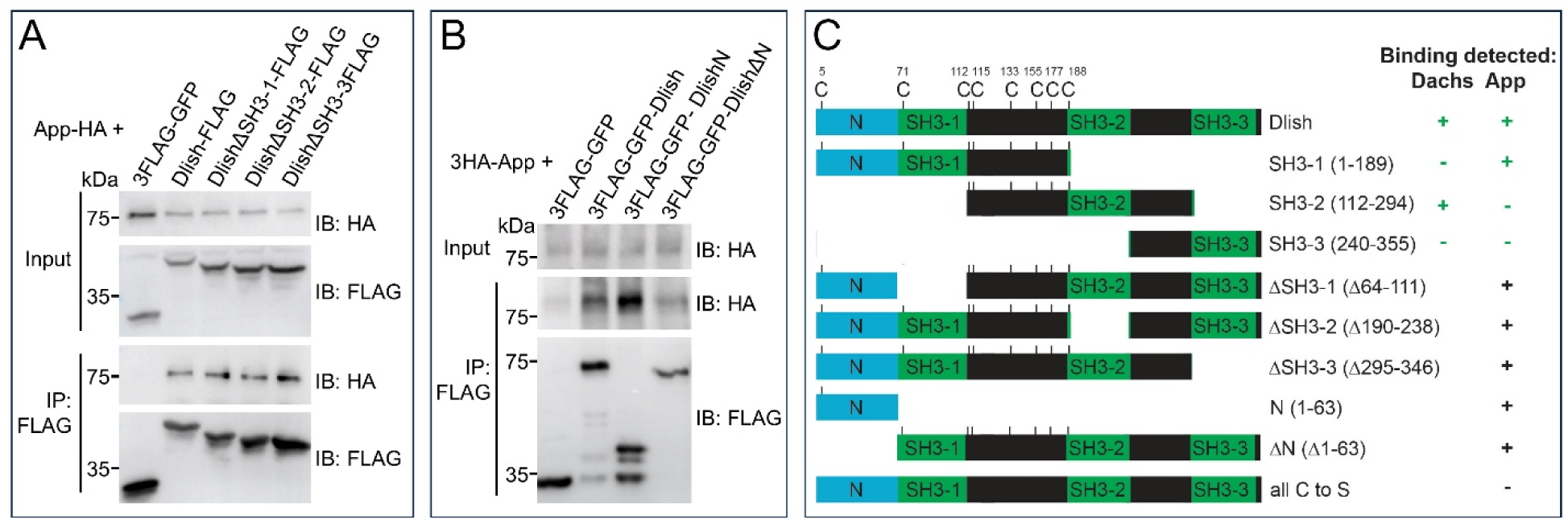
Role of Dlish N terminus mediating Dlish-App binding in S2R+ cells. A. In vitro assay of IP between co-expressed App-HA and Dlish-FLAG lacking specific SH3 domains. B. In vitro assay of IP between co-expressed 3HA-App and 3FLAG-GFP-Dlish, 3FLAG-GFP-DlishN, and 3FLAG-GFP-DlishΔN. C. Summary of constructs and co-IP results; co-IP results in green are from ref^15^.

The second SH3 domain of Dlish is necessary and sufficient for binding Dachs in vitro; it is also necessary for normal cortical stabilization of Dachs in vivo^15,16^. DlishΔSH3-2 should thus lose Dachs binding but retain App binding, and will also retain all of the potential palmitoylated cysteines (Fig. 8C). As noted above, when expressed in vivo DlishΔSH3-2 concentrated in the subapical cortex, even after knockdown of Dachs or loss of endogenous Dlish (Fig. 3H-J). However, because it cannot bind Dachs it does not detectably improve subapical Dachs in a *dlish* null mutant background (Fig. 9A,A’) or alter growth in a wild type background (Fig.8B).

**Figure 9.**
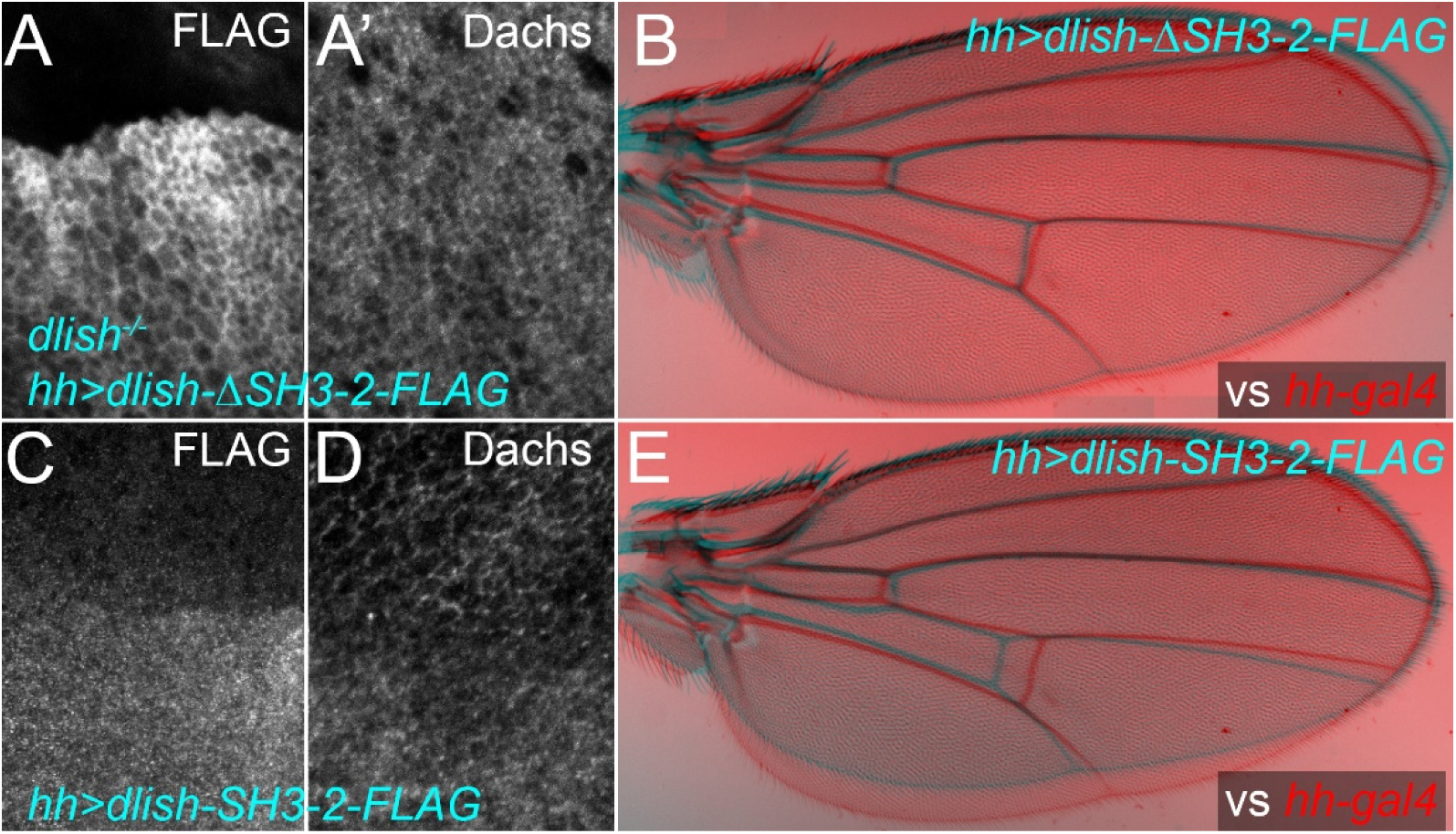
In vivo localization and activity of Dlish constructs with stronger and weaker App binding. A,A’. Localization of posterior, *hh-gal4*-driven *UAS-dlishΔSH3-2-FLAG* in *dlish^B1601^* homozygous wing disc and its effects on anti-Dachs. B. Comparison of wing growth in *hh-gal4* control (red) and *hh-gal4 UAS-dlishΔSH3-2-FLAG* (cyan). C,D. Localization of posterior, *hh-gal4*-driven *UAS-dlish-SH3-2-FLAG* in wing disc (C) and its effects on anti-Dachs (D). E,E’. Comparison of wing growth in *hh-gal4* control (red) and *hh-gal4 UAS-dlishSH3-2-FLAG* (cyan).

In contrast a Dlish construct retaining SH3-2 (Dlish-SH3-2) and all of the potential palmitoylation targets except C5 and C71, but lacking the NT, SH3-1 and SH3-3 domains (Fig. 8C), should retain Dachs binding but lack App binding and thus palmitoylation. As expected, when overexpressed Dlish-SH3-2 concentrated in the cytoplasm rather than the cell cortex and reduced cortical and increased cytoplasmic Dachs, likely through binding Dachs and recruiting it to the cytoplasm (Fig. 9C,D), and resulted in undergrowth like that seen after *dachs* knockdown (Fig. 9E).

## Discussion

We have shown here that interactions between the multi-SH3 domain protein Dlish, the atypical myosin Dachs, and the palmitoyltransferase App play overlapping but distinct roles mediating the effects of the ICDs of Fat and Ds on components of the Hippo pathway. We discuss below the implications of our findings for different models of Fat and Ds activity.

### App and palmitoylated Dlish

First, our results confirm that, in normal cells, direct binding between cortical Dlish and Dachs helps increase the levels of Dachs at the subapical cortex and reduce Dachs levels in the cytoplasm. Our results also strongly support our hypothesis that the cortical localization and activity of Dlish depends on its palmitoylation by App. The activity and cortical localization of Dlish is blocked by removal of its potential palmitoylation sites. We think it more likely that palmitoylation anchors Dlish to the cell membrane, rather than to a specific protein partner, as the loss of Dlish palmitoylation can be counteracted by adding a prenylation site distant from the normal palmitoylation sites, and thus less likely to cooperate with other protein interaction motifs. We also provide evidence that normal Dlish localization and activity depend on Dlish’s ability to bind App and confirm that loss of App or Dlish have similar effects on Dachs. We note, however, that while loss of *fat* cannot increase Dachs levels in *dlish* null mutants, *fat* loss still weakly increases Dachs in *app* null mutants^33,40^; either Dlish retains weak activity after loss of palmitoylation or its palmitoylation is not wholly dependent on App.

### Dlish as a mediator of Fat’s effects on Dachs

Dlish’s effects, however, go beyond the cortical localization of Dachs: we found that cells lacking *dlish* can retain residual Dachs in the subapical cortex but this Dachs is no longer sensitive to the accumulation of junctional Ds or the loss of Fat. Similarly, Matakatsu and Fehon found that, without Dlish, Dachs is no longer sensitive to simultaneous loss of Fat and Ds^33^. The insensitivity of the residual cortical Dachs to *fat* loss in a *dlish* null strongly suggests that Dlish not only helps localize Dachs, but that Dlish is, or complexes Dachs with, a mediator of the Fat ICD.

The binding data suggests several possible models of Fat activity. In vitro, Dlish, App, and Dachs can all bind to each other and each independently bind the Fat ICD; Dlish and Fat as well bind to other possible effectors^15,16,23,33,40^. We here ruled out one possible effector; while the scaffolding protein Ex directly binds both Dlish and the Fat ICD^23,35^, loss of Ex does not elicit or block the increased cortical Dlish caused by *fat* knockdown.

### Palmitoylation inhibition models

We previously proposed a model that could account for the Dlish-dependent activity of the Fat ICD: that the Fat ICD binds to App and reduces either App’s levels and/or its access to its normal targets. In *fat* mutants more App would be free to palmitoylate Dlish, increasing cortical accumulation of the Dlish/Dachs complex^2,15^ (Fig. 10A). Changes in App levels might mediate the effects of the Fat’s ICD as loss of both *fat* and *ds* increases cortical App, albeit not as strongly as it increases Dlish and Dachs^33^.

**Figure 10.**
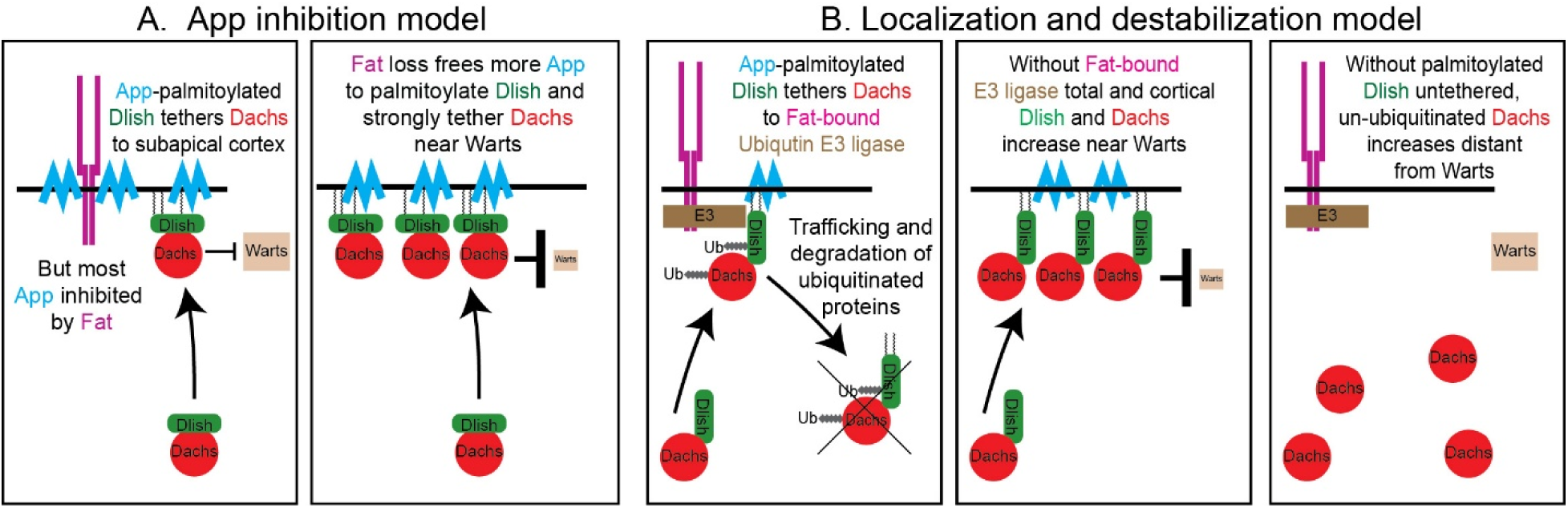
- Models of Fat ICD signal transduction

However, our evidence argues that App is present in excess and that changes in App levels alone are unlikely to account for Fat’s activity: very large reductions in App levels have no effect on wing development. Conversely, strong overexpression of App does not increase, but instead slightly reduces, both growth and cortical Dlish^17,33^.

It is still possible that App activity requires an unknown, limiting co-factor that is itself inhibited by the Fat ICD. Another version of the inhibition model is that the Fat ICD sequesters Dlish away from App, again reducing its palmitoylation. But given the high levels of non-junctional and cytoplasmic Dlish and App, especially after overexpression, this also seems unlikely. Thus, the results to date suggest that, in the context of the wing imaginal disc, Dlish palmitoylation by App is largely permissive, not instructive.

### Localization and destabilization model

Another problem with the palmitoylation inhibition models is that while they might explain the effects of Fat and Dlish on the cortical localization of Dachs, they do not address the effects on total Dachs levels, which increase 2-3-fold in the absence of either Fat or Dlish^15,33^. A non-exclusive alternative is that Fat ICD organizes a complex that destabilizes Dlish and Dachs in the subapical cortex, and that palmitoylated Dlish not only recruits Dachs to the cortex but also couples Dachs to this destabilizing complex (Fig. 10B). Loss of Fat would stabilize cortical Dlish-bound Dachs; loss of Dlish would stabilize Dachs but most of that Dachs would be lost from the cortex. Overexpression of Dlish would have two opposing effects, recruiting Dachs to the cortex but also coupling Dachs to the destabilizing complex, consistent with the observed weak increase in Dachs^15^.

This “localization and destabilization” model is supported by work on the Fbxl7 F-box E3 ubiquitin ligase: the Fat ICD binds to and increases cortical levels of Fbxl7, *fbxl7* mutants increase cortical levels of Dachs and Dlish (as well as Fat and Ds), and Dlish can bind Fbxl7^15,48,49^. Fbxl7 activity is not, however, sufficient to explain all of the *fat* or *dlish* mutant phenotype, as the effects of losing *fbxl7* on cortical Dlish/Dachs are weaker than loss of *fat*, and total Dachs levels are not detectably increased^15,48^. Studies also differed on whether Dachs is ubiquitinated by Fbxl7^48,49^. Nonetheless, this hypothesis is attractive since Dlish can regulate ubiquitination in a different context; in vitro Dlish binds both Ex and the Slimb F-box E3 ligase and Dlish increases the Slimb-based ubiquitination of Ex^23^. In the context of Fat and Dlish, the hypomorphic *slimb^1^* mutation can also weakly increase cortical Dlish and Dachs^15^, so Fbxl7 may be only one of several E3 ligases involved.

### Contrasting roles for Dlish and Dachs

So far our Fat ICD models have focused on App and Dlish as mediators, but the effects of Dlish and Dachs on each other are in some sense reciprocal: they form a complex, both bind the Fat and Ds ICDs, overexpression of one can increase cortical levels of the other, and mutation of either blocks the overgrowth of *fat* mutations^14-16,33^. However, while loss of Dlish both mislocalizes Dachs and increases its total levels, we found that knockdown or loss of Dachs very strongly decreases Dlish levels in both the cortex and cytoplasm, enough so that it is unclear if the small amount of remaining endogenous Dlish is or is not sensitive to the loss of Fat. Moreover, Dachs is unnecessary for Dlish localization: overexpression of Dlish was sufficient to overcome the absence of Dachs, and that Dlish still preferentially localized to the subapical cell cortex in the absence of Dachs. Dlish is needed to localize Dachs near its cortical targets, explaining why overexpresssed Dachs only weakly elicits overgrowth in *dlish* mutants^15^. Dachs is not needed to localize Dlish, explaining why overexpressed Dlish induces strong overgrowth in *dachs* mutants^23^.

We do not know how Dachs protects Dlish, but if it acted locally it could mediate some portion of Fat’s effect on Dlish, for example if the Fat ICD reduced Dach’s ability to bind or otherwise protect Dlish. And any Dlish-mediated increase in cortical Dachs would elicit local positive feedback by further protecting Dlish, which in turn would increase the recruitment of Dachs to the cortex. Our working hypothesis invokes Dlish as an adaptor for E3 ubiquitin ligases that can target either Dlish or Dachs. Without Dachs Dlish is ubiquitinated; when Dachs binds Dlish it protects it, either by blocking ligase access or by acting as a more favored target. However, our tests reducing the activity two E3 ubiquitin ligases known to affect Dlish levels did not detectably increase Dlish levels after Dachs knockdown.

#### Regulation by the Ds ICD-

In vivo Dachs accumulates at cell junctions with high Ds, and as Dachs binds the Ds ICD in vitro, the in vivo effect is thought to be mediated by simple binding^13,33^. We were therefore surprised that Ds overexpression did not detectably affect Dachs in *dlish* mutants, despite the residual Dachs present in the subapical cortex. It is possible that robust binding between Dachs and Ds requires Dlish; however, more complex models are possible that utilize variants on the regulatory models proposed for the Fat ICD.

## Methods

### Fly strains

*From stock centers, labs, or previous studies*

*UAS-GFP*; *P(Gal4)hh^Gal4^*/*TM6,Tb* = *hh-gal4*^57^

*hh-gal4 ban3-GFP/TM6,Tb* (^58^)

*y^1^ w*; P{w^+mW.hs^=en2.4-GAL4}e16E = en-gal4* (BDSC 30564)

*P{w^+mC^=UAS-Dcr-2.D}1, w^1118^; en-gal4 P{w^+mC^=UAS-2xEGFP}AH2* (BDCS 25752)

*P{w^+mC^=UAS-Dcr-2.D}1, w^1118^; hh-gal4/TM6,Tb*

*ap^md544^* (*ap-gal4*) *UAS-gapGFP*/*CyO* (from BDSC 3051 and BDSC 4522)

*UAS-dachs-V5* (^15^)

*dachs-GFP* (^59^)

*fat^fd^ FRT^40A^ dlish^B1601^*/*CyO,Tb*

*NM^31E^ FRT^40A^ dlish^Y003^*; *hh-gal4 UAS-Flp*/*CyO-TM6,Tb* (^23^)

*UAS-ds* (on III; ^42^)

*hh-gal4 UAS-ds/TM6,Tb FRT^42D^ dlish^B160123^*

*FRT^42D^ dlish^B1601^; hh-gal4 /TM3,Sb*

*FRT^42D^ dlish^B1601^; hh-gal4 /SM5a-TM6B*

*FRT^42D^ dlish^B1601^; UAS-ds/SM5a-TM6B*

*ex^e1^ FRT^40A^/CyO* (BDSC 44249)

*y^1^ v^1^; P{y[+t7.7] v[+t1.8]=TRiP.JF03245}attP2 (UAS-fat-RNAi;* BDSC 29556)

*UAS-d-RNAi (P{KK111834}VIE-260B*; VDRC v102504)

*en-gal4 UAS-d-RNAi/CyO*

*d^GC13^ FRT^40A^*/*CyO,Tb* (BDSC 28289)

*d^210^ pr; hh-gal4/CyO-TM6B* (from BDSC 6389)

*UAS-fbxl7-RNAi* (*y^1^ v^1^; P{y^+t7.7^ v^+t1.8^=TRiP.JF01515}attP2*; BDSC 31065)

*UAS-elgi-CS* (^41^), provided by K. Irvine

*FRT^2A^ app^12-3^/TM6,Tb* (^40^; provided by H. Matakatsu)

*y w hsFlp*; *FRT^2A^ ubi-GFP* (from BDSC 1626)

*w^1118^; Df(3L)BSC730/TM6C, Sb^1^ cu^1^* (BDSC 26828)

*UAS-app-RNAi-VDRC* (*w^1118^; P{GD9293*}; VDRC v32863)

*UAS-app-RNAi-TRIP (y^1^ w^1^; P{TRiP.GL00181}attP2;* BDSC 35280)

*app^YFP^ hh-gal4/TM6,Tb* (^33^), provided by H. Matakatsu

*UAS-dcr2*; *app^3xV5^ hh-gal4*/*TM6,Tb* (^33^), provided by H. Matakatsu

*UAS-dlish-FLAG* (^15^)

*hh-gal4 UAS-dlish-FLAG/TM6,Tb*

*P{ w^+mC^=sqh-EYFP-Golgi}3* (BDSC 7193)

*P{ w^+mC^=sqh-EYFP-ER}3* (BDSC 7195)

*y^1^ w^1118^; P{w^+mC^=UASp-GFP-mCherry-Atg8a}2* (BDSC 37749)

*y^1^ M{w^+mC^=myc-wts-V5}ZH-2A w\** (BDSC 56809)

*y^1^ w*; Mi{PT-GFSTF.1}Cad99C^MI08439-GFSTF.1^* (BDSC 60268)

#### New for this study

The stocks below are insertions at the attP-9A site on chromosome III using y*^1^ w* P{nanos-phiC31\int.NLS}X; PBac{y+-attP-9A}VK00027)* (BDSC 35569)

*UAS-dΔSH3bd-3-FLAG*

*FRT^42D^ dlish^B1601^*; *UAS-dlish*Δ*SH3-2-FLAG*/*TM3,Sb*

*UAS-dlish-all C to S-FLAG UAS-dlish-FLAG-CAAX*

*hh-gal4 UAS-dlish-FLAG-CAAX*

*UAS-dlish-all C to S-FLAG-CAAX*

*UAS-dlish-SH3-2-FLAG*

To induce hsFLP expression for creating homozygous FRT-based mitotic recombination clones, larvae were treated in a 37°C water bath for 1.5h.

### Staining and microscopy

Immunostaining, microscopy and image processing of imaginal discs and adult wings was as previously described^15,23,60^, with the following additions. Most discs stained with anti-Dachs and/or our original anti-Dlish^15^ were fixed at room temperature for 5m in PBS with 2% formaldehyde. Mouse anti-FLAG M2 (Sigma) was used at 1:500-2,500, rabbit anti-GFP (MBL International JM-3999-100) at 1:500, mouse anti-V5 (Invitrogen 46-0705) at 1:500, rabbit anti-Ds^61^; (provided by David Strutt) was used at 1:1,000. To rule out cross-reactive secondaries for Dlish-CAAX-FLAG or anti-Ds colocalization studies, discs were first stained for the critical primary antigen and fluorescent secondary before being stained for anti-FLAG or anti-Ds. A new guinea pig anti-Ex generated for this study and used at 1:5,000, and a new rabbit anti-Dlish was used at 1:5000 for discs in Fig. 6 (antiserum generation shown below).

#### LysoTracker

*-* Live discs were incubated in a 1:500 dilution in PBS of a LysoTracker Deep Red (Thermo Fisher Scientific L12492) stock solution for 30-40m and washed in PBS. Live discs were mounted in PBS under coverslips with spacers made from the paper backing of Parafilm and sealed with petroleum jelly, or discs were fixed 30m at 4°C in 4% formaldehyde in PBS and stained with anti-GFP and anti-FLAG.

### Generation of new anti-Ex and anti-Dlish

A pGEX-4T construct encoding Ex amino acids 145–947 and a pET28b(+) construct encoding full length Dlish were transformed into *E. coli* Rosetta (DE3) competent cells. GST-Ex was induced with 1 mM IPTG at 37°C for 4h and purified using glutathione-Sepharose beads (Ge Healthcare, 17-0756-01). His-Dlish was induced with 100 mM IPTG at 25°C for 12h and purified using HisPur Ni-NTA Superflow Agarose (Thermo 25214). Proteins were resolved by SDS–PAGE, visualized by Coomassie Brilliant Blue staining, and the corresponding gel bands cut and used for guineapig (Ex) or rabbit (Dlish) immunization by ABclonal Technology. Antisera were affinity-purified using Protein G beads.

### DNA constructs

The following plasmids have been described previously^15^)(^23^: paw-Gal4, pUAST-attB-Dlish-FLAG, pUAST-attB-Dlish-HA, pUAST-3FLAG-Ajuba, pUAST-attB-App-FLAG, pUAST-Dachs-FLAG, pUAST-attB-Dco-FLAG, pUAST-FtΔECD-FLAG, pUAST-attB-DsICD-FLAG, pUAST-attB-Fbxl7-FLAG, pUAST-attB-Dachs ΔSH3bd-3-FLAG, pGEX2T-Dlish.

The following new plasmids were built using In-Fusion HD Cloning Plus (Clontech 638909) or ClonExpress Ultra One Step Cloning Kit (Vazyme C115):pUAST-attB-Dlish all C to S-FLAG, pUAST-attB-Dlish (aa 5, 71, 112, 115, 155, 177, 188) C to S-FLAG, pUAST-attB-Dlish (aa 5, 71, 112, 115, 177, 188) C to S-FLAG, pUAST-attB-Dlish (aa 5) C to S-FLAG, pUAST-attB-Dlish (aa 133) C to S-FLAG, pUAST-3HA-Dlish all C to S-attB, pUAST-attB-App-HA, pUAST-3FLAG-GFP, pUAST-attB-Dlish-ΔSH3-1-FLAG (Δaa64-111), pUAST-attB-Dlish-ΔSH3-2-FLAG (Δaa190-238), pUAST-attB-Dlish-ΔSH3-3-FLAG (Δaa295-346), pUAST-3FLAG-Dlish, pUAST-3FLAG-DlishN, pUASt-3FLAG-DlishN, pUAST-3FLAG-GFP-Dlish-N (aa1-63), pUAST-3FLAG-GFP-Dlish-ΔN (aa64-355), pMBP-App(aa1-349), and pMBP-App(aa350-694).

### In vitro analyses

Culture of Drosophila S2R+ cells, transfection, harvesting, protein extraction, co-IP, and SDS-PAGE were as previously described^15^.

#### MBP pull-down assay

*-* GST-tagged Dlish was prepared from *E. coli* as previously (Zhang et al., 2016). MBP, MBP-App(aa1-349), and MBP-App(aa350-694) were expressed in BL21(DE3)pLysS *E. coli* competent cells (Promega, L1195) and purified with Amylose Resin (NEB, E8021S) in accordance with the manufacturer’s recommendations. The purity and quantity of proteins were analyzed by SDS-PAGE and Coomassie staining and western blotting. 4 μg purified MBP, MBP-App(aa1-349) and MBP-App(aa350-694) protein were mixed with 0.5 μg purified GST-Dlish in binding buffer (1X PBS, 0.1%NP40, 10% Glycerol, 0.5 mM DTT, protease inhibitor) with protease inhibitor and incubated at 4°C for 2h. 20 μl of washed Amylose Resin was added for an additional 1h, washed and then boiled for 8m with 25 μl 2× SDS/protein sample buffer prior to loading for SDS-PAGE.

#### Western blotting and imaging

- As described previously^15^ using the following antisera: Mouse anti-MBP (CW Biotech, CW0288M) at 1:6000, Mouse anti-GST (Santa Cruz, sc-138) at 1:6000, Rabbit anti-FLAG (Sigma-Aldrich F7425) at 1:7000, Rabbit anti-FLAG (EasyBio BE2005) at 1:10000, Rabbit anti-HA (EasyBio BE2008) at 1:9000. Protein was visualized either with a) HRP secondaries, the chemoluminescent ECL kit and either autoradiograph film or the Fuasion Solo Vilber imaging system, or b) IRDye®-secondaries (Li-Co Biosciences) and the Li-COR Odyssey imaging system.

#### Palmitoylation assays

- The Acyl-Biotin-Exchange (ABE) method was as described previously^15^ except that proteins were immunoprecipitated with mouse anti-FLAG agarose beads (Sigma-Aldrich Cat#A2220) and the blot in Fig. 5A was stained with both IRDye® 680RD Streptavidin (1:5000) (LI-COR Biosciences Cat# 926–68079) and with anti rabbit-FLAG primary (1:7000) (Sigma-Aldrich Cat# F7425 Lot# RRID:AB_439687) and IRDye® 800CW Goat anti-Rabbit IgG (H + L) secondary (1:10000) (LI-COR Biosciences Cat# 925–32211).

For the click chemistry assay, 24h after transfection S2R+ cells were starved in serum-free Schneider’s medium for 1h to deplete endogenous lipids, incubated 6h with 100 μM alkyne palmitic acid (15-hexadecynoic acid, Cayman Chemical). and then lysed in ice-cold lysis buffer (50 mM Tris-HCl, pH 7.4, 150 mM NaCl, 1% Triton X-100) supplemented with protease inhibitors 500 μg of total protein lysate was conjugated with Biotin-PEG4-Azide using the Click-iT™ Protein Reaction Buffer Kit (Thermo Fisher Scientific) according to the manufacturer’s instructions, and then immunoprecipitated using anti-FLAG Magnetic Beads (Thermo Fisher Scientific). To verify the thioester linkage of palmitoylation, bead-bound proteins were divided into two equal aliquots and incubated with either 1 M hydroxylamine (Sigma-Aldrich, pH 7.0) or 1 M Tris-HCl (pH 7.0) for 1 h at room temperature. Following extensive washing, the proteins were eluted by boiling in 2× SDS loading buffer, followed by SDS-PAGE and western blotting. Palmitoylated Dlish was detected with IRDye® 800CW Streptavidin (LI-COR Biosciences) and total FLAG-Dlish with anti-FLAG and an IRDye® 680RD secondary.

### Informatics

We used FlyBase (release FB2023_05) to find information on gene and RNA sequences, mapping of RNAi targets, Drosophila stocks, and publications.

## Acknowledgements

This work was supported by grants from the NIH (R01-GM124377 and R01-GM151072; SSB), the Natural Science Foundation of Beijing, China (No. 5252010; XW), and the National Natural Science Foundation of China (No. 32270528; YZ).

We thank Drs Matakatsu and Fehon, Dr. Irvine, the Vienna Drosophila Resource Center (VDRC), the TRiP at Harvard Medical School (NIH/NIGMS R01-GM084947), and the Bloomington Drosophila Stock Center (NIH P40OD018537) for fly stocks used in this study, and Drs D. Stutt and H. McNeill for antisera.

## Author contributions

X.W., Y.Z., and S.S.B. conceived the project, designed the experiments, analyzed and interpreted the data, and prepared the manuscript. X.W., Y.A., J.Z., X.Y., and S.S.B. performed the experiments.

---

**Figure S1.**
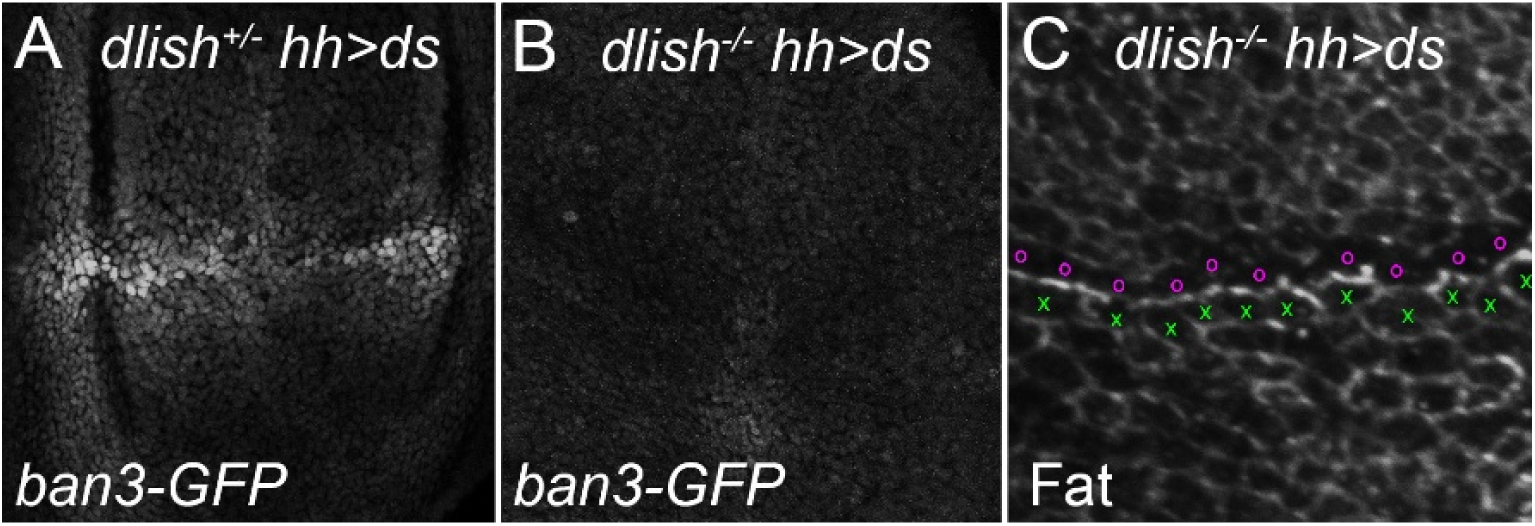
Effect of *dlish* loss on the activation of the Yki target *ban3-GFP* by boundaries of Ds overexpression in wing discs. A,B Expression of *ban3-GFP* after posterior, *hh-gal4-driven* expression of *UAS-ds* in wild type (A) and *dlish^B1601^* homozygous (B) backgrounds. C. Depletion of anti-Fat staining from non-boundary cell faces (left and right) of cells just anterior to boundary (marked by magenta o) in *dlish^B1601^* homozygote with *hh-gal4-driven* expression of *UAS-ds* (cells at anterior limit marked by green x).

**Figure S2.**
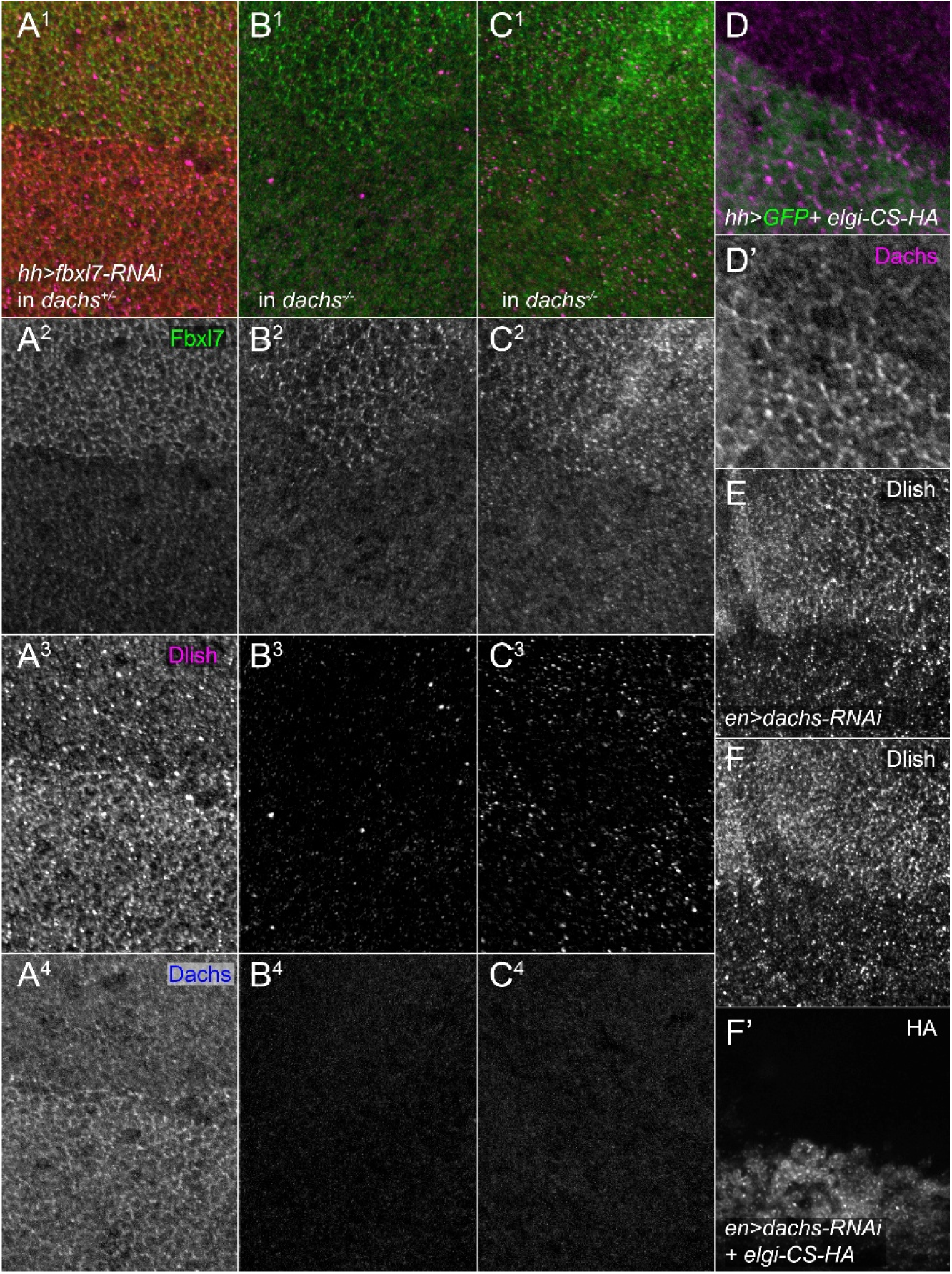
The effect of reducing of Fbxl7 or Elgi activity on the loss of Dlish caused by *dachs* loss or knockdown. A^1^-C^4^. Posterior, *hh-gal4*-driven expression of *UAS-fbxl7-RNAi* and its effect on anti-Fblx7 staining (green in A^1^,B^1^,C^1^ and grey in A^2^,B^2^,C^2^), anti-Dlish staining (magenta in A^1^-C^1^, grey in A^3^-C^3^), and anti-Dachs staining (blue in A^1^-C^1^, grey in A^4^-C^4^) in +/ *d^210^* (A1-A4) or *d^GC13^/d^210^* wing discs (B^3^,C^3^) identified by the absence of anti-Dachs staining (B1^-B4^,C1-C4). D,D’. Anti-Dachs staining (magenta in D, grey in D’) after posterior, *hh-gal4*-driven expression of the dominant negative *UAS-elgi-CS-HA*, identified by co-expression of *UAS-GFP* (green in D). E-F’. Anti-Dlish staining (E,F) after posterior, *en*-gal4-driven expression of U*AS-dachs-RNAi*, without (E) or with{F) *UAS-elgi-CS-HA*, identified by anti-HA staining (F’). Discs being compared (those in A^1^-C^4^, and those in E and F) were fixed and stained in the same well and imaged and processed with the same settings.

**Fig. S3.**
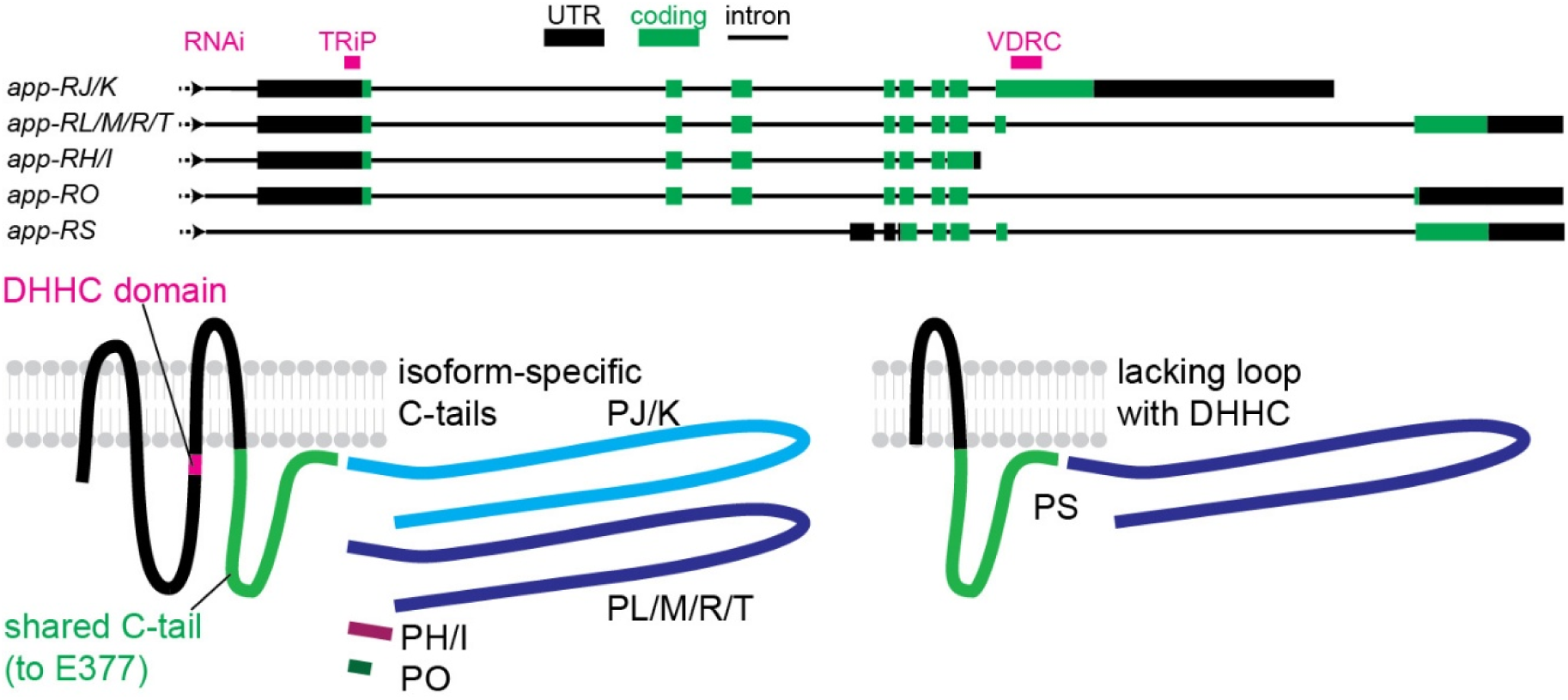
Top: predicted alternative splicing of *app* transcript and targeting by the TRiP and VDRC *UAS-app-RNAi*’s. Bottom: predicted protein isoforms. Data from ref{Matakatsu, 2008 #757} and Flybase^62^.

**Figure S4.**
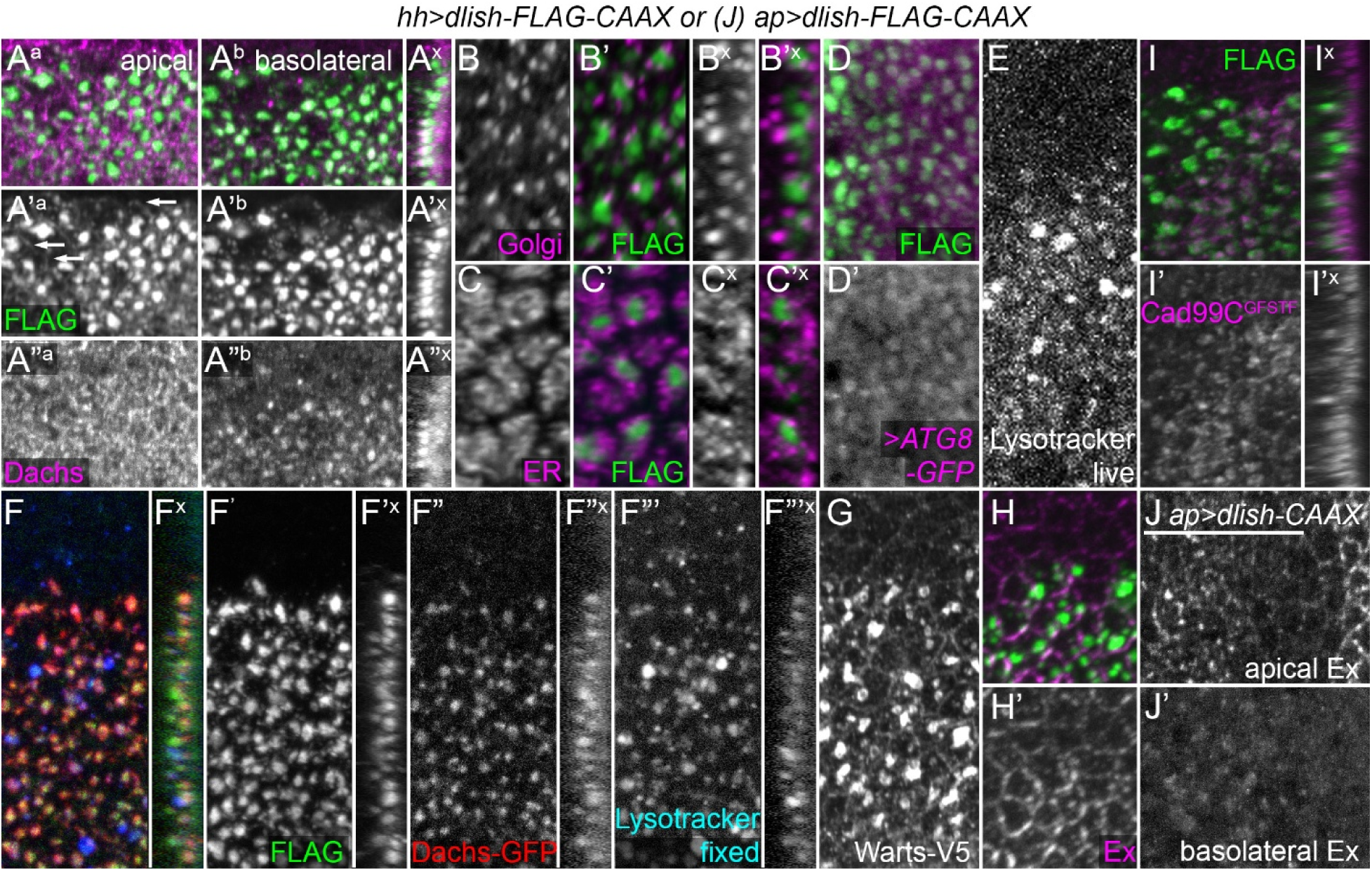
A-I. Colocalization of large anti-FLAG staining aggregates (green) after posterior, *hh-gal4*-driven expression of *UAS -dlish-FLAG-CAAX* with autolysosomal markers and other proteins. Panels marked with superscript x are cross-sections with apical to the right. A^a^-A”^x^. Anti-FLAG staining in membranes (A’^a^, arrows) and aggregates; the latter partially overlap with anti-Dachs especially in slightly more basolateral sections (A^b^-A”^b^, cross-sections in A^x^-A”^x^). B-C’^x^. Little overlap between FLAG aggregates and cell compartment markers EYFP-Golgi (B) or EYFP-ER (C), both expressed via *sqh*-derived regulatory sequences^63^. D,D’. Overlap between FLAG aggregates and *UAS*-driven ATG8-GFP. E. Posterior induction of Lysotracker staining of live discs. F-F”’^x^. Partial overlap between FLAG aggregates, Dachs-GFP, and Lysotracker staining of live discs imaged after fixation. G. Induction of Warts-V5 aggregates. H,H’. No accumulation of anti-Ex staining in FLAG aggregates. I,I’. Partial overlap of the GFP-tagged Cad99C^GFSTF^ protein trap with FLAG aggregates. J,J’. Weak accumulation of basolateral Ex aggregates after dorsal (left), *ap-gal4-*driven expression of *UAS-dlish-FLAG-CAAX*.

**Figure S5.**
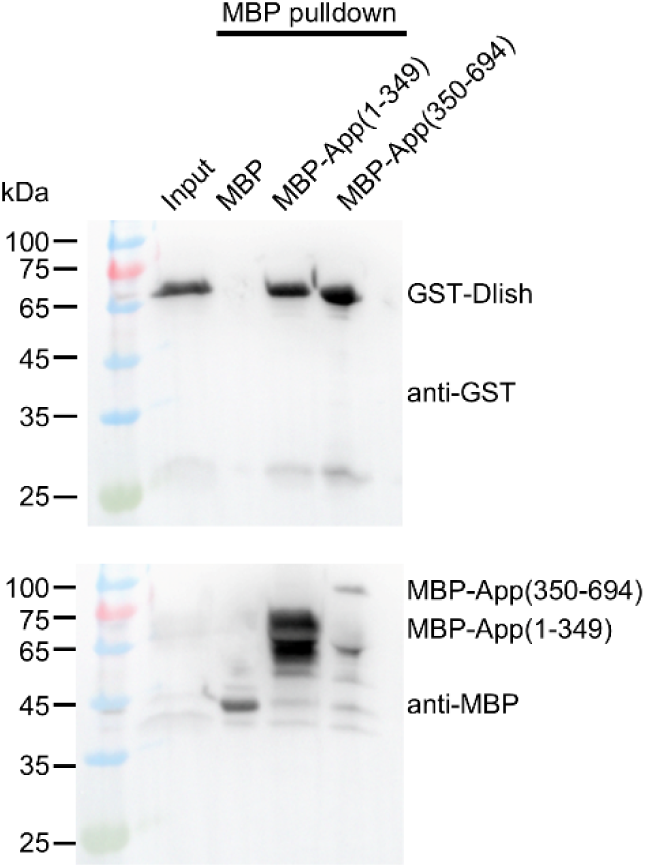
Pulldown of bacterially-produced GST-Dlish by bacterially produced MBP-App(1-349) or MBP-App(350-694).

